# Conserved neural population geometry supports behavioral generalization

**DOI:** 10.1101/2024.10.24.620127

**Authors:** Sara A Solla, John F Disterhoft, Hannah S Wirtshafter

**Author notes:** Correspondence and requests for materials should be addressed to Hannah S Wirtshafter,. Lead contact (HSW).

## Abstract

Learned behaviors often generalize across contexts even though neural representations can vary substantially between environments. How the brain preserves task-relevant information across these changing neural representations remains unclear. This problem is particularly evident in the hippocampus because hippocampal spatial representations reorganize across environments even when learned behaviors generalize successfully. Here, we recorded hippocampal neuronal activity using calcium imaging while rats performed a conditioning task in two distinct environments. Despite remapping of spatial representations, the low-dimensional organization of task related population activity was conserved across contexts, and the temporal relationships among neural population states during task execution were preserved. Strikingly, this relational organization was conserved not only across contexts for individual animals, but also across animals, despite differences in the underlying neural populations and individual experiences. Moreover, this conserved organization was sufficient to support transfer of task decoding across animals, demonstrating that task information could generalize between independently learned neural representations without shared neuron identities. These findings identify a conserved population level neural organization through which the hippocampus can support behavioral generalization despite contextual remapping and suggest that shared organizational neural mechanisms and structures may underlie behavioral generalization across individuals.

## Introduction

Learned behaviors often generalize across contexts even though neural representations of different contexts can vary substantially. How the brain preserves task-relevant information across these changing neural representations remains a central question in neuroscience.

This problem is particularly evident in the hippocampus (HPC) because hippocampal spatial representations^1–5^ reorganize across environments, a phenomenon known as remapping^6–10^; surprisingly, learned behaviors generalize successfully despite this remapping^6–8^. Because hippocampal activity also encodes nonspatial task related variables^11–16^, a major and unresolved question arises: what aspects of task related population activity are preserved when contextual representations reorganize? A further open question is whether different animals solve this problem using shared population level strategies, or through idiosyncratic and varied neural solutions.

To address these questions, we used *in* vivo calcium imaging to record hippocampal CA1 activity while rats performed an HPC-dependent conditioning task, trace eyeblink conditioning (tEBC), which is rapidly generalized between spatial contexts^17,18^. Animals learned the task in one environment and were then tested in a different environment, one that triggered the remapping of spatial representations. We compared representations of spatial location and task structure across environments and across animals.

Although spatial representations remapped across environments, task related population activity preserved a stable low-dimensional geometry during behavioral generalization, as represented by the relationships among task states. Strikingly, this relational organization was conserved not only across contexts for individual animals, but also across animals despite differences in neural populations and experiences. Moreover, this conserved organization was sufficient to support transfer of task decoding across animals without shared neuron identities.

Together, these findings identify a conserved population level organization of hippocampal task representations that persists across both contextual remapping and individual variability. More broadly, these results suggest that neural systems can preserve a relational task structure even as context-dependent representations reorganize.

## Results

We trained five freely moving rats in a conditioning task across two distinct environments, labeled A and B; animals were familiarized only with environment A before conditioning training began (Fig. 1a, Fig. S1). Environment A was an unscented rectangular enclosure with white lighting, whereas environment B was a scented ovoidal enclosure with red lighting (Fig. 1b, Fig. S1). Both environments were placed in the same position relative to the external room cues (see Methods). During daily training and testing sessions, we recorded hippocampal CA1 neuronal activity using miniature microscopes^19^ (Miniscopes) and the calcium indicator GCaMP8m. Analyses used either deconvolved calcium events or calcium traces, as specified (see Methods).

**Figure 1.**
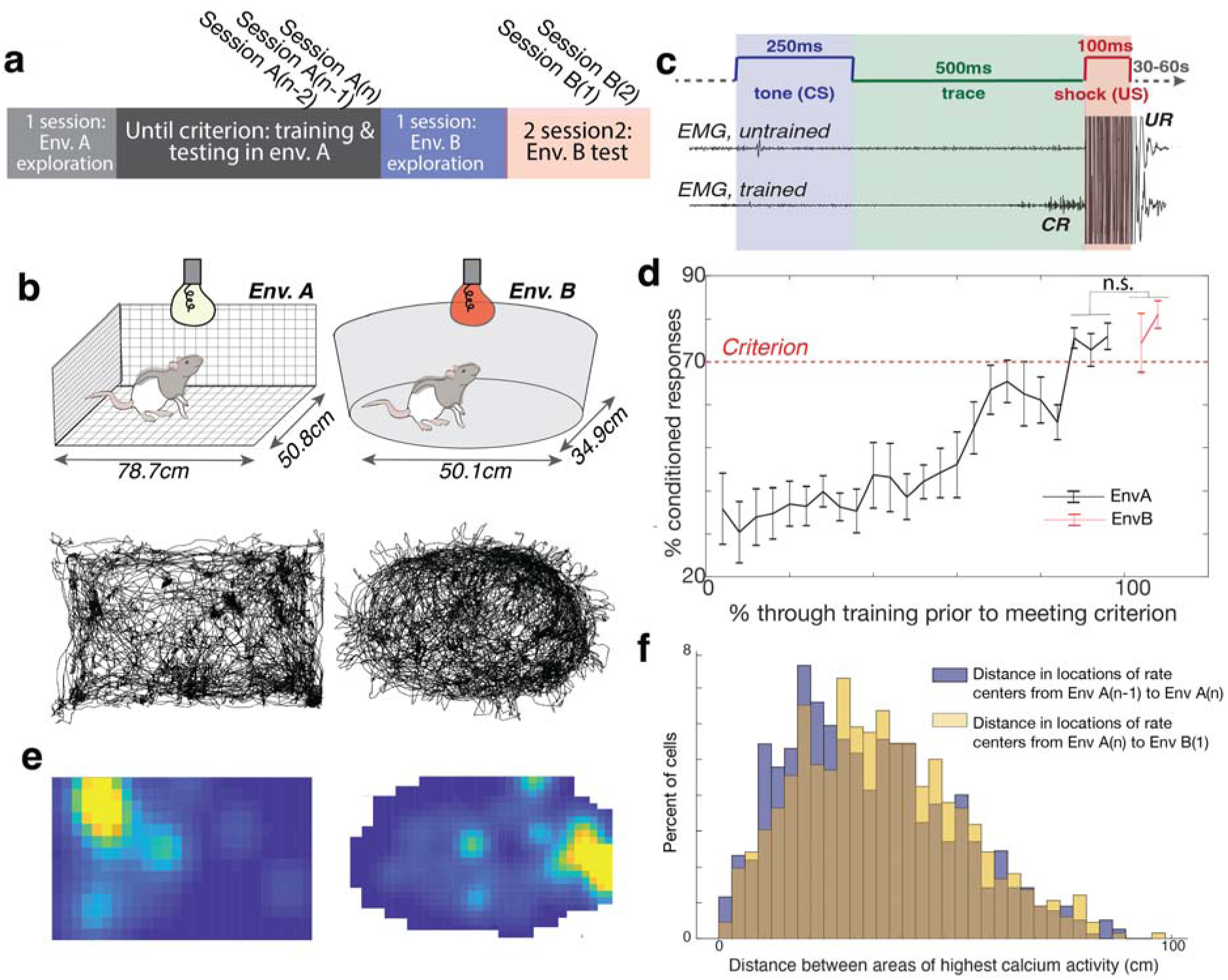
Conditioning is maintained while cells remap between environment A and environment B. **a.** Animals explored environment A for one session before undergoing trace eyeblink conditioning for *n* sessions until they reached criterion in environment A (See Methods). Animals were then allowed one session of exploration in environment B, followed by two tEBC test sessions in environment B, indicated as B(1) and B(2). **b.** Top: Schematics of environments A and B. Environment A is a rectangular enclosure with wire walls, floor, and ceiling, lit with white light, and unscented. Environment B is ovoidal with solid white floors and walls, without a ceiling, lit with red light, and scented with clove oil. Both environments provided distal cues visible from the top and sides. Bottom: Representative animal trajectory for single sessions in environments A and B, respectively, extracted using DeepLabCut. **c.** Trace eyeblink conditioning (tEBC) paradigm. A 250ms tone (conditioned stimulus, CS) was followed by a 500ms trace interval and a 100ms eyelid shock (unconditioned stimulus, US). Eyelid activity was recorded using an EMG electrode implanted above the eye. Untrained animals only blinked in response to the US (unconditioned response, UR); trained animals began blinking during the trace interval after the CS and before the US (conditioned response, CR). **d.** Performance of all rats (n=5) in the tEBC task. Animals learned tEBC while freely moving in environment A and successfully transferred this learning to environment B. The dotted line indicates the performance criterion. No significant difference was found between performance in the criterion sessions in environment A (mean 74.75 ± 6.49%) and the testing sessions in environment B (mean 77.70 ± 11.68%; two-tailed t-test, t(24) = -0.83, p > 0.05). Error bars represent SEM. **e.** Map of the activity of a single example cell in environments A and B. Yellow indicates the highest activity. This cell exhibited remapping between environments; it shows different rate centers relative to external cues in the two contexts. **f.** Distribution of distances between rate centers when comparing sessions A(n-1) and A(n) versus comparing sessions A(n) and B(1). Rate centers shifted significantly more when the animal was moved to environment B, compared to shifts between two consecutive sessions in environment A (Wilcoxon rank sum test, p = 0.005; two-sided t-test, t(1430) = -2.5, p = 0.01; two-sample Kolmogorov-Smirnov test, p = 3.0 × 10^-3^). The brown shaded area represents the overlap between the two histograms.

### Animals generalize a conditioning task across environments despite hippocampal remapping

Freely moving rats (Fig. 1b) were trained on trace eyeblink conditioning (tEBC), a hippocampus-dependent associative learning task^20–23^. In this paradigm, a 250ms auditory conditioned stimulus (CS; tone) is followed by a 500ms trace interval and then a 100ms unconditioned stimulus (US; eyelid shock) (Fig. 1c). As training progressed, animals developed conditioned blink responses (CRs) to the tone and were considered to have learned the task upon reaching behavioral criterion (70% correct for three days, occurred in a mean of 20 ± 4.2 training sessions; see Methods, Fig. 1d). After reaching criterion in environment A, rats were transferred to environment B for one day of exploration followed by two days of testing on the tEBC task without additional training.

Comparative analysis revealed no significant difference in behavioral performance between the final three training sessions in environment A and the two test sessions in environment B (mean %CRs: 74.75 ± 6.49 in environment A versus 77.70 ± 11.68 in environment B; two-sided t-test, t(24) = −0.83, p > 0.05; Fig. 1d), indicating successful transfer of the learned behavior to a new environment.

Consistent with hippocampal remapping^1,8,24,25^, individual CA1 neurons exhibited distinct spatial representations in environments A and B (Fig. 1e). For each neuron, we identified the spatial location of peak calcium event rate (‘rate centers’) during non-conditioning periods^16^ and quantified the stability of spatial representations by measuring the distance between rate centers across sessions and across environments. Rate centers shifted significantly more between environments than between sessions in the same environment (two-sided t-test, t(1430) = −2.5, p = 0.01; Fig. 1f), indicating a reorganization of spatial representations across contexts. In contrast, shifts of rate centers within environments were significantly smaller than expected by chance; in shuffled controls, rate centers were randomly reassigned across neurons within each session (Fig. S2). Spatial decoding analyses yielded similar results (Fig. S3). These findings indicate that animals generalized the learned behavior despite robust hippocampal remapping across environments.

### Task related latent representations generalize across environments

We next asked whether hippocampal population activity contained a low-dimensional representation of the conditioning task whose organization generalized across environments. Calcium imaging data from session A(n) were divided into two labeled epochs corresponding to the CS/trace and US/post-US periods; we refer to this representation as CSUS2 to emphasize the segmentation of the task into two time bins (Fig. 2a). We used a nonlinear method for dimensionality reduction, the CEBRA^26^ algorithm, to obtain low-dimensional latent embeddings of the population activity during task trials.

**Figure 2.**
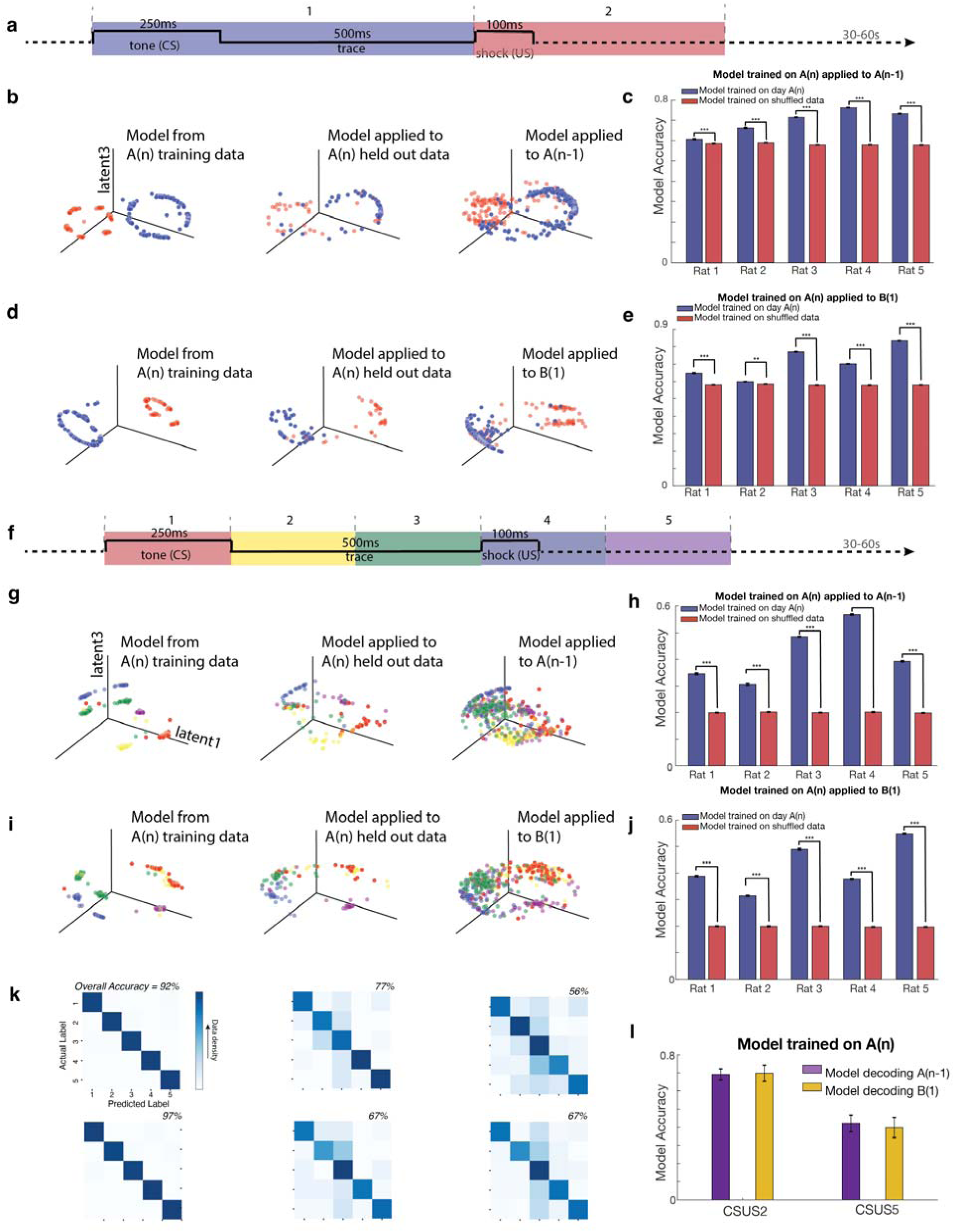
Models trained in environment A generalize across environments to decode task epochs and fine temporal structure. **a.** Schematic of the trace eyeblink conditioning (tEBC) task and the CSUS2 temporal segmentation. The conditioning period was divided into two epochs: a CS/trace period (blue; 250ms tone + 500ms trace interval) and a US/post-US period (red; 100ms shock + post-US period). Intertrial intervals ranged from 30s to 60s. **b.** Example CEBRA embeddings for CSUS2 decoding using neurons shared between sessions A(n) and A(n−1) (Rat 3 example from one representative CEBRA run of embedding and decoding). Each point represents neural population activity at a specific epoch, projected into the latent space learned using data from the A(n) session. Left: model trained on task data from 75% of trials in session A(n) (25% of data held out as test data). Middle: latent representation of held out A(n) data (25%) projected into the latent space learned in one CEBRA run using the complementary 75% of session A(n) task data. Right: latent representation of data from session A(n−1) projected into the latent space shown on the left, learned in one CEBRA run using 75% of session A(n) task data. Note that the separation between the two epochs was preserved across sessions within the same environment. **c.** Decoding accuracy, averaged across 500 CEBRA embedding runs for each rat, for CSUS2 models trained on session A(n) and applied to session A(n−1). For all five rats, models trained with data from session A(n) decoded CS vs US in data from session A(n−1) significantly better than if decoding data with shuffled epoch labels (all two-sided t-tests: Rat 1: t(998) = 8.0, p = 4.9 × 10^-15^; Rat 2: t(998) = 35.8, p = 3.6 × 10^-181^; Rat 3: t(998) = 83.5, p = 0; Rat 4: t(998) = 80.1, p = 0; Rat 5: t(998) = 61.8, p = 0). Black error bars indicate SEM; significance denoted as ***p < 10^-10^. **d.** Example CEBRA embeddings for CSUS2 decoding using neurons shared between sessions A(n) and B(1) (Rat 3 example from one representative CEBRA run of embedding and decoding). Each point represents neural population activity at a specific epoch, projected into the latent space learned using data from the A(n) session. Left: model trained on task data from 75% of trials in session A(n). Middle: latent representation of held out A(n) data (25%) projected into the latent space learned in one CEBRA run using the complementary 75% of session A(n) task data. Right: latent representation of data from session B(1) projected into the latent space shown on the left, learned in one CEBRA run using 75% of session A(n) task data. Note that the separation between the two epochs generalized across environments. **e.** Decoding accuracy, averaged across 500 CEBRA embedding runs for each rat, for CSUS2 models trained on session A(n) and applied to session B(1). For all five rats, models trained with data from session A(n) decoded CS vs US in data from session B(1) significantly better than if decoding data with shuffled epoch labels (all two-sided t-tests: Rat 1: t(998) = 16.8, p = 7.6 × 10^-56^; Rat 2: t(998) = 3.0, p = 3.2 × 10^-3^; Rat 3: t(998) = 75.3, p = 0; Rat 4: t(998) = 63.7, p = 0; Rat 5: t(998) = 90.6, p = 0). Black error bars represent SEM; **p < 10^-3^, ***p < 10^-10^. **f.** Schematic of the CSUS5 temporal segmentation used for fine temporal decoding analyses of the tEBC task. The conditioning period was divided into five sequential temporal epochs: one CS epoch (red), two trace epochs (yellow, green), and two epochs spanning the US and post-US periods (blue, purple). **g.** Example CEBRA embeddings for CSUS5 decoding using neurons shared between sessions A(n) and A(n−1) (Rat 3 example from one representative CEBRA run of embedding and decoding). Each point represents neural population activity at a specific epoch, projected into the latent space learned using data from the A(n) session. Left: model trained on task data from 75% of trials in session A(n). Middle: latent representation of held out A(n) data (25%) projected into the latent space learned in one CEBRA run using the complementary 75% of session A(n) task data. Right: latent representation of data from session A(n−1) projected into the latent space shown on the left, learned in one CEBRA run using 75% of session A(n) task data. Note that fine temporal structure within the task period was preserved across sessions within the same environment. **h.** Decoding accuracy, averaged across 500 CEBRA embedding runs for each rat, for CSUS5 models trained on session A(n) and applied to session A(n−1). For all five rats, these models significantly outperformed the decoding of data with shuffled epoch labels (all two-sided t-tests: Rat 1: t(998) = 34.8, p = 2.0 × 10^-174^; Rat 2: t(998) = 33.6, p = 7.8 × 10^-166^; Rat 3: t(998) = 122.6, p = 0; Rat 4: t(998) = 118.5, p = 0; Rat 5: t(998) = 62.6, p = 0). Black error bars indicate SEM; significance denoted as ***p < 10^-10^. **i.** Example CEBRA embeddings for CSUS5 decoding using neurons shared between sessions A(n) and B(1) (Rat 3 example from one representative CEBRA run of embedding and decoding). Each point represents neural population activity at a specific epoch, projected into the latent space learned using data from the A(n) session. Left: model trained on task data from 75% of trials in session A(n). Middle: latent representation of held out A(n) data (25%) projected into the latent space learned in one CEBRA run using the complementary 75% of session A(n) task data. Right: latent representation of data from session B(1) projected into the latent space shown on the left, learned in one CEBRA run using 75% of session A(n) task data. Note that fine temporal structure within the conditioning period generalized across environments. **j.** Decoding accuracy, averaged across 500 CEBRA embedding runs for each rat, for CSUS5 models trained on session A(n) and applied to session B(1). For all five rats, these models outperformed the decoding of data with shuffled epoch labels, indicating that fine temporal task structure generalized across environments (all two-sided t-tests: Rat 1: t(998) = 55.1, p = 4.9 × 10^-305; Rat 2: t(998) = 22.8, p = 7.0 × 10^-93; Rat 3: t(998) = 71.4, p = 0; Rat 4: t(998) = 62.6, p = 0; Rat 5: t(998) = 106.2, p = 0). Black error bars indicate SEM; significance denoted as ***p < 10^-10. **k.** Confusion matrices for the classification of latent data onto the five epochs in CSUS5, based on CEBRA decoding of the five CSUS5 temporal bins shown in panels g (top row) and i (bottom row) (Rat 3 example from one representative CEBRA run of embedding and decoding). Columns show model performance on A(n) training data, held out A(n) data, and transfer decoding on A(n−1) (top row) or B(1) (bottom row). Darker diagonal values indicate higher decoding accuracy. Overall decoding accuracy, given by the sum of diagonal elements divided by the sum of all elements, is shown for each matrix. **l.** Decoding accuracy, averaged across 500 CEBRA embedding runs for each rat, for data from sessions A(n−1) and B(1) using models trained on 75% of data from session A(n) with labels provided by CSUS2 and CSUS5 temporal segmentations. For both CSUS2 and CSUS5, models trained on data from session A(n) decoded session B(1) with accuracy comparable to that for decoding session A(n−1) (CSUS2: two-sided t-test, t(8) = −0.13, p > 0.05; CSUS5: t(8) = 0.32, p > 0.05). Black error bars represent SEM.

The activity during each CSUS2 task trial resulted in two clusters of points in the latent space, one for activity during the CS epoch and one for activity during the US epoch. The CEBRA algorithm provides an embedding of these points using two latent dimensions, displayed on the surface of a three-dimensional sphere. The epoch labels identified structure in the latent space: the embeddings of data from environment A formed distinct and separable latent trajectories corresponding to the two task epochs. Decoders trained to separate CS points from US points using data from session A(n) reliably generalized to session A(n−1) (Fig. 2b,c). For all animals, decoding performance significantly exceeded that of shuffled controls generated by permuting the task labels corresponding to data points in latent space (all two-sided t-tests, p < 0.001), indicating that the latent representation of the task was stable across sessions within the same environment.

We next asked whether this latent organization generalized across contextual remapping. Activity from session B(1), the first tEBC testing in the novel environment, exhibited a similar structure to that of A(n) when projected into the latent space learned from environment A (Fig. 2d). Additionally, decoders trained using only data from session A(n) successfully distinguished the two task epochs in B(1), significantly outperforming shuffled controls generated by permuting the correspondence between latent space points and epoch labels for each animal (Fig. 2e; all two-sided t-tests, p < 0.001). These results demonstrate that hippocampal population activity contains a temporally structured latent representation of the conditioning task that generalizes across environments.

To determine whether finer temporal structure within the conditioning period was similarly preserved, we next used a representation of the task divided into five epochs (CSUS5): one CS epoch, two trace epochs, and two epochs spanning the US and post-US periods (Fig. 2f). At this finer time resolution, the CEBRA embeddings of data from environment A showed smooth progressions through latent space, with neighboring task epochs occupying nearby positions and more distant epochs occupying positions further apart (Fig. 2g). Decoders trained to separate the five time bins in CSUS5 using data from session A(n) reliably generalized to session A(n−1) (Fig. 2g), significantly outperforming shuffled controls generated by permuting the correspondence between latent space points and epoch labels for each animal (Fig. 2h; all two-sided t-tests, p < 0.001). Confusion matrices showed strong decoding along the diagonal, indicating preservation of identity of neural states corresponding to each task epoch across sessions within the same environment (Fig. 2k).

We next asked whether this finer latent structure generalized across contextual remapping. Activity from session B(1) exhibited a similar progression of temporal epochs when embedded using the model learned from session A(n) (Fig. 2i). Decoders trained using data from session A(n) accurately decoded activity from session B(1), again significantly outperforming shuffled controls for each animal (Fig. 2j; all two-sided t-tests, p < 0.001). Confusion matrices showed strong decoding along the diagonal, indicating preservation of the identity of neural states corresponding to each task epoch across environments (Fig. 2k).

Remarkably, for both CSUS2 and CSUS5, decoding accuracy in B(1) did not differ from decoding accuracy in A(n−1) (two-sided t-tests, both p > 0.05; Fig. 2l), indicating that cross-context transfer was comparable to within-context transfer. The transfer of task information was not affected by the remapping associated with the change of context.

Although animals overall showed behavioral transfer to environment B, transfer strength varied somewhat across individuals. One animal (Rat 2) showed weaker behavioral transfer during session B(1), the first testing session in the new environment; this was accompanied by reduced transfer of the latent representation. However, both behavioral performance and decoding accuracy improved substantially by session B(2), where CSUS2 decoding accuracy reached almost 70%. This significant improvement over day B(1) (two-sided t-test, p<0.001, Fig. S4) was in line with the stabilization of the transferred task representation with increased exposure to the novel environment.

In contrast to the observed transfer of task representation across environments, models trained to decode spatial position in environment A did not generalize reliably to environment B, as expected given the remapping of spatial representations (Fig. S3). Thus, generalization across environments was selective for task related temporal structure and did not imply a general preservation of hippocampal population activity across environments.

### Task related latent geometry is preserved across environments

While the decoding analyses shown above demonstrate that task related information generalizes across environments, successful decoding alone does not establish whether the underlying organization of neural states is preserved. Therefore, we next examined the structure of latent task representations using several complementary analyses, including consistency scores, geometry-preservation metrics, and trajectory-alignment measures. Together, these analyses assess whether task representations retain a common relational organization across environments despite hippocampal remapping.

To directly test whether tEBC representations share a conserved population geometry across environments, we compared the structure of task related activity in the CEBRA embedding space. Task representations were highly similar across environments A and B, with CEBRA consistency scores (a measure of similarity between independently trained latent embeddings, see Methods) significantly exceeding controls generated by randomly shuffling time bin labels prior to model training across multiple embedding dimensions. This consistency was robust for both coarse (CSUS2) and fine (CSUS5) temporal segmentations of the task (Fig. 3a). All five rats showed significantly greater correlations in task structure across environments than shuffled controls; this effect remained significant across embeddings based on up to 10 latent dimensions (Fig. 3a-b).

**Figure 3.**
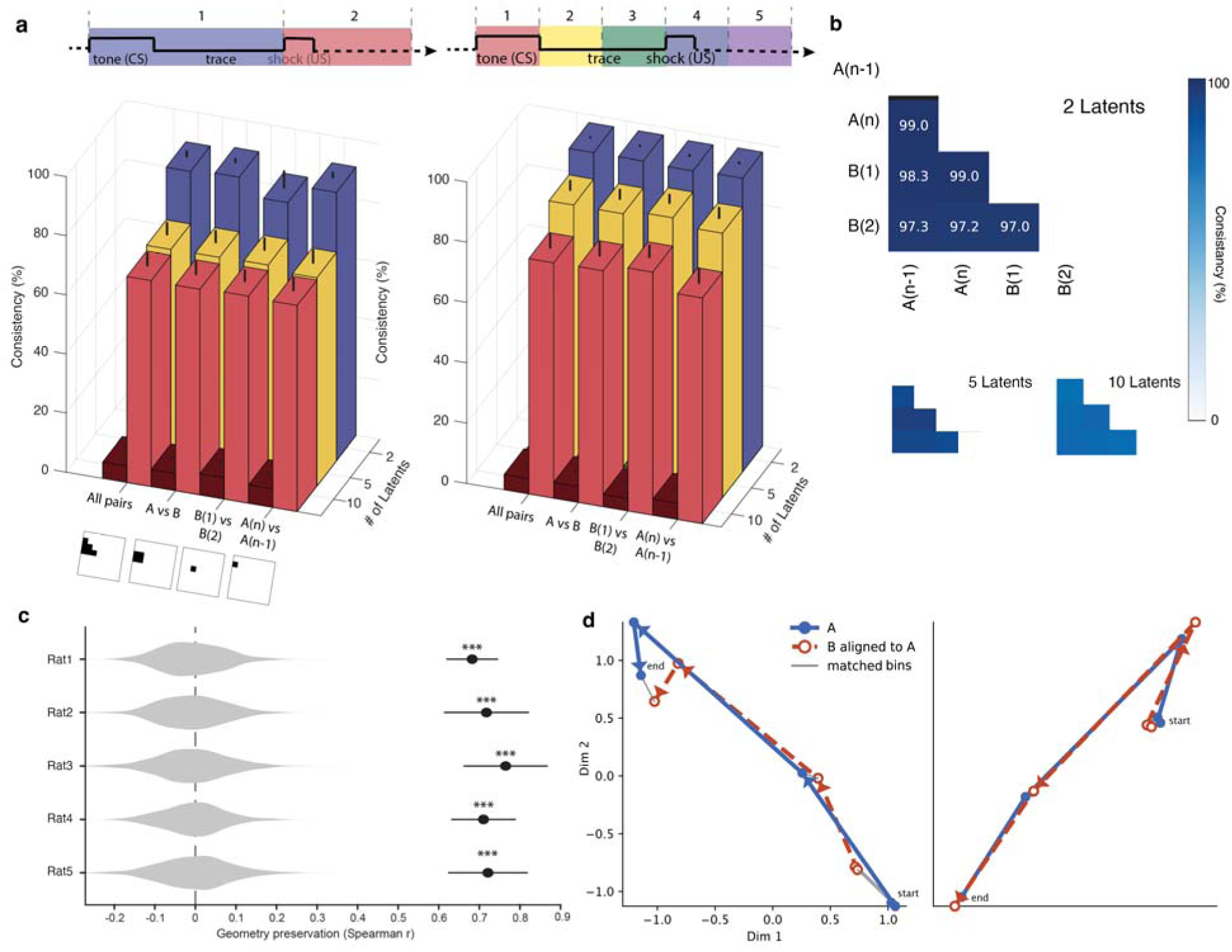
Conserved population geometry of tEBC latent task representations across environments. a. Top: Schematics of the CSUS2 and CSUS5 temporal segmentations of the tEBC task. Bottom left: CEBRA consistency scores for CSUS2 latent representations with 2, 5, and 10 latent dimensions. Colored bars show consistency scores for actual data, while adjacent darker bars show scores obtained after randomly shuffling time bin labels. The datasets included in each comparison are indicated below each of the four sets of bars. “All pairs” includes all six pairwise comparisons among A(n−1), A(n), B(1), and B(2): A(n−1) vs A(n), A(n−1) vs B(1), A(n−1) vs B(2), A(n) vs B(1), A(n) vs B(2), and B(1) vs B(2). “A–B pairs” includes only the four cross-environment comparisons: A(n−1) vs B(1), A(n−1) vs B(2), A(n) vs B(1), and A(n) vs B(2). For aggregated comparisons, consistency was computed separately for each pairwise comparison and then averaged. Each bar shows mean ± SEM across rats. Consistency scores significantly exceeded shuffled controls across all latent dimensions (all two-sided t-tests: 2 latents, p < 1 × 10^⁻^□; 5 latents, p < 1 × 10^⁻^□; 10 latents, p < 1 × 10^⁻^□). Bottom right: Same analysis for CSUS5. All five rats showed significantly greater consistency scores than those for shuffled controls. Significant differences between actual and shuffled data were observed for all latent dimensions (all two-sided t-tests: 2 latents, p < 1 × 10^⁻¹²^; 5 latents, p < 1 × 10^⁻^□; 10 latents, p < 1 × 10^⁻^□). b. Example consistency matrices from Rat 3 for CSUS5 using 2, 5, and 10 latent dimensions. Consistency scores remained high across environments for actual data but not for models trained on shuffled time bin labels. Matrix components involving models trained on shuffled data are omitted for clarity; their consistency scores were uniformly low (<12%). c. Geometry preservation across contexts. For each rat, CSUS5 task state centroids were computed for each time bin by averaging latent coordinates for that time bin across all trials, for sessions A(n) and B(1) separately. Pairwise Euclidean distances were then computed among the five time bin centroids within each environment, yielding 10 unique pairwise distances per session. Geometry preservation across environments was quantified as the Spearman correlation between these arrays of distances for A(n) and B(1). Violin plots show control distributions generated by randomly permuting CSUS5 time bin labels in B(1) before recomputing geometry preservation scores. Blue circles indicate mean values of geometry preservation for the real data; horizontal lines denote ±SEM across CEBRA model runs. Geometry preservation exceeded shuffled controls in all rats (one-sided permutation tests, 500 permutations); in all rats, no shuffled value exceeded the observed mean value (p = 0.002). d. Aligned task trajectories for two example animals (Rat 2, left; Rat 5, right). Environment B(1) embeddings (red) are shown after orthogonal Procrustes alignment to the corresponding environment A(n) embeddings (blue) for the same rat. Each trajectory represents the latent trajectory averaged across all trials for that session. Points correspond to the five time bins (1–5), with “start” and “end” indicating bin #1 and bin #5, respectively. For each rat, gray lines connect corresponding bins across environments. Because the orientation of the latent space is arbitrary, trajectories for different rats are shown in independent coordinate frames.

We next asked whether the relational organization of intermediate task related neural population states was preserved across environments. Latent trajectories during tEBC followed similar paths in sessions A(n) and B(1), preserving both the temporal ordering and the relative arrangement of task states across successive time bins (Fig. 3c). Geometry preservation, quantified from the pairwise distances among task states in latent space, significantly exceeded shuffled controls generated by permuting time bin labels in environment B (Fig. 3c; in all rats, no shuffled value exceeded the observed value, p = 0.002, 500 permutations; see Methods). Notably, although one animal exhibited comparatively weaker decoding transfer across environments, its latent geometry remained largely preserved, indicating that geometry preservation and decoder transfer are distinct metrics for quantifying the generalization of task representations across environments.

To visualize this shared geometric structure directly, latent trajectories for each environment were aligned using the orthogonal Procrustes algorithm^27,28^, which finds the rotation (and possible reflection) that best matches corresponding task states while preserving the geometry of the latent space (Fig. 3d). After alignment, corresponding task states occupied similar relative positions across environments. Consistent with this observation, for all rats, orthogonal Procrustes error was significantly lower than expected from controls obtained by training models on shuffled time bin labels (all p < 0.002; Fig. S5). This demonstrates that task trajectories were substantially more alignable across contexts than expected by chance. These results indicate that contextual generalization is associated with the preservation of the relational organization among task related neural population states.

Together, the results in Fig. 3 indicate that hippocampal representations of the tEBC task are organized in a stable, low-dimensional population geometry that preserves the temporal organization of the task despite contextual remapping.

### Task related latent geometry is conserved across both animals and environments

As described above, latent task representations were highly similar across environments for each individual animal. We next asked whether task representations also shared a common organization across animals. To address this question, we trained CEBRA models for each animal using time bin labels and quantified the CEBRA consistency scores between models for different animals.

Latent task representations exhibited high cross-animal consistency relative to controls generated from models trained on shuffled time bin labels. This effect was observed both for the coarse task segmentation (CSUS2) (Fig. 4a) and for the finer temporal segmentation of the task (CSUS5) (Fig. 4b).

**Figure 4.**
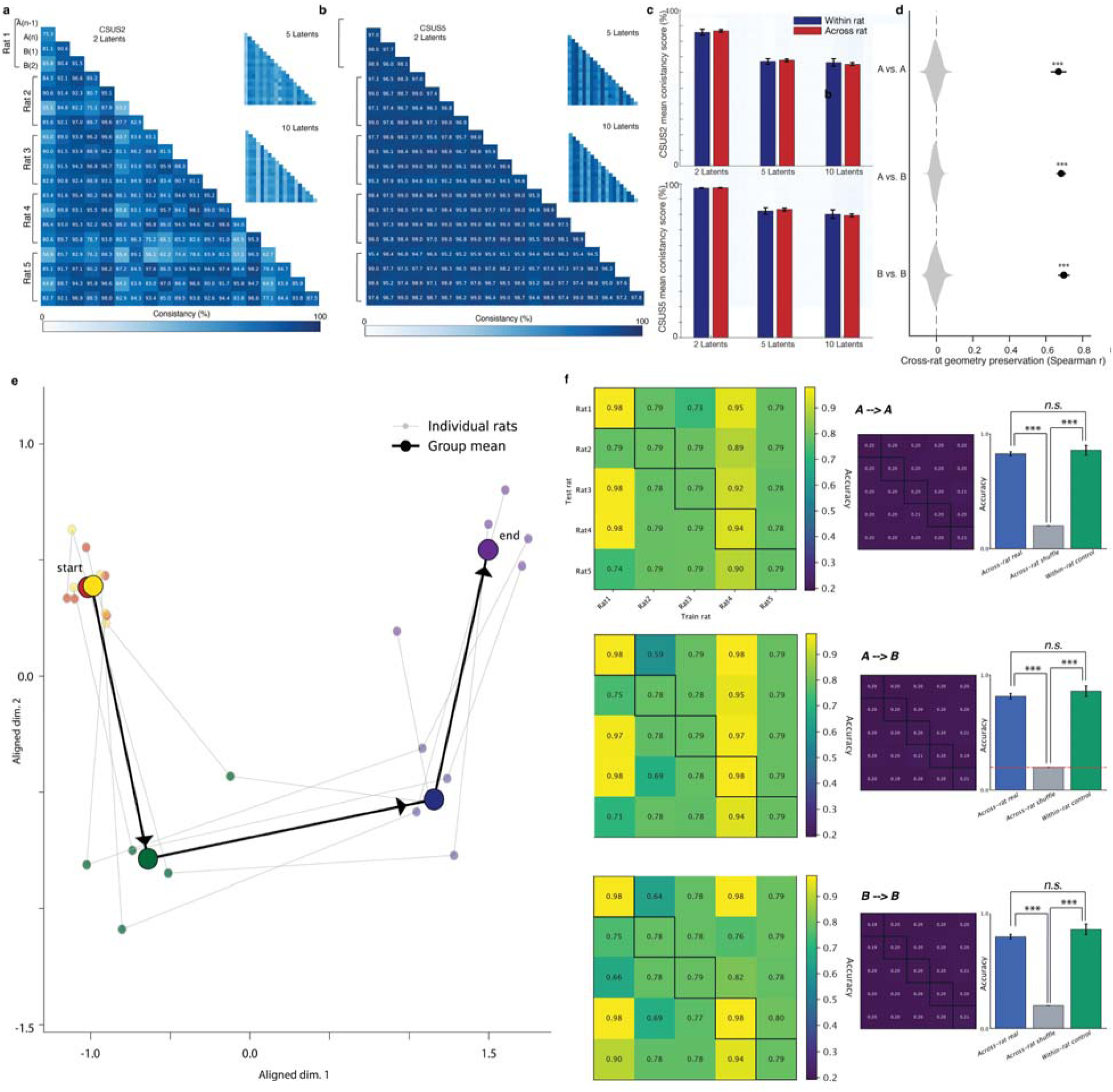
Conserved population geometry of tEBC latent task representations across animals. a. CEBRA consistency scores across animals and across sessions for CSUS2 latent representations. Heat maps show pairwise consistency scores across rats and across sessions A(n−1), A(n), B(1), and B(2) for 2 latent dimensions (left), and 5 and 10 latent dimensions (right). Additional matrix components involving models trained on shuffled data were uniformly low (<12%) and are omitted for clarity. High consistency scores were observed across animals for all latent dimensionalities. b. Same analysis as in panel a, but for CSUS5 representations divided into five time bins. c. Bar graphs compare across animals CEBRA consistency scores (blue) with consistency scores across environments for a given animal (red). For all latent dimensionalities (2, 5, and 10), consistency scores across animals did not differ significantly from same animal consistency scores (all p > 0.05, two-sided t-tests). Bars show mean ± SEM for CSUS2 (top) and CSUS5 (bottom). d. Geometry preservation across rats was quantified as the correlation between CSUS5 matrices of pairwise distances between centroids of latent representations for each time bin. Comparisons were performed separately for A(n) versus A(n), B(1) versus B(1), and A(n) versus B(1). For each comparison, geometry preservation was computed for each rat pair and CEBRA model run, and then averaged across the ten rat pairs. Shuffled null distributions were generated by randomly permuting time bin labels in one member of each comparison before recomputing geometry preservation scores. Circles indicate mean geometry preservation scores, and horizontal lines denote ±SEM. e. Task trajectories aligned across animals, using orthogonal Procrustes. CSUS5 task trajectories for each rat after orthogonal Procrustes alignment are shown as light gray lines in a shared two-dimensional latent space. Each trajectory represents the average across sessions A(n) and B(1) for that rat. The mean trajectory over all five rats is shown in black. Points correspond to the five time bins (1–5), with “start” and “end” indicating bin #1 and bin #5, respectively. f. Conserved latent geometry of the conditioning task supports decoding transfer across animals. Task epoch identity for CSUS5 was decoded across rats using independently trained CEBRA embeddings, without matching neurons across animals. The top row shows decoding across rats for sessions A(n)→A(n), the middle row shows decoding for sessions A(n)→B(1), and the bottom row shows decoding for sessions B(1)→B(1). For each comparison, latent spaces were aligned using orthogonal Procrustes alignment, and a logistic-regression decoder trained on one rat (train rat) was evaluated on another rat (test rat). Left: Heatmaps show the decoding accuracy for each ordered pair, with ‘train rat’ on the x-axis and ‘test rat’ on the y-axis. These heatmaps show the decoding accuracy obtained from real labels; the diagonal entries show controls where the same rat is used for training and testing. Middle: Heatmaps show decoding accuracy for shuffled controls obtained by permuting the time bin labels for the train rat before the decoder is trained. Right: Bar plots summarize mean decoding accuracy across all off-diagonal pairs involving a test rat different from the train rat (cross-rat real), the corresponding controls with shuffled labels on the train rat (cross-rat shuffle), and diagonal controls for decoding on the same rat as training (within-rat control). Mean off-diagonal decoding accuracies were 0.826 for A(n)→A(n), 0.819 for A(n)→B(1), and 0.798 for B(1)→B(1). Cross-rat decoding was significantly greater than shuffled controls for all comparison types (paired two-sided t-tests, A→A: t(19)=35.81, p=6.61×10^⁻¹^□; A→B: t(19)=25.51, p=3.67×10^⁻¹^□; B→B: t(19)=29.48, p=2.50×10^⁻¹^□) and did not differ significantly from within-rat decoding performance (two-sided t-tests, A→A: p=0.54; A→B: p=0.44; B→B: p=0.25). Error bars show mean ± SEM. ***p < 0.001; n.s., not significant.

We then asked whether the similarity captured by the consistency analysis reflected preservation of latent geometry across animals. Remarkably, geometry preservation remained high even when comparing animals that experienced different environments (A vs. B). Across all latent dimensionalities examined, between 2 and 10, geometric similarity between different animals was indistinguishable from the similarity observed across repeated sessions, for both CSUS2 and CSUS5 representations (all p > 0.05, two-sided t-tests; Fig. 4c). These findings suggest that hippocampal task representations share a common geometric organization across individuals, one that is maintained despite the substantial variability introduced by both contextual remapping and differences between animals.

We next asked whether this similarity reflected preservation of the distance relationships among task states in latent space. To test this, we repeated the analysis from Fig. 3c–d across animals, using mean task trajectories averaged across trials in sessions A(n) and B(1), for each rat. Geometry preservation was significantly greater than chance for all comparison types (Fig. 4d), demonstrating that the relational organization among successive task states during tEBC was conserved across animals.

After alignment, trajectories for different animals preserved both the temporal ordering and relative arrangement of neural population states across CSUS5 bins (Fig. 4e). The aligned trajectories revealed a conserved progression of neural population states across animals. In line with this observation, the residual orthogonal Procrustes disparity after alignment was substantially lower than expected from shuffled controls for all comparisons (all permutation p < 0.002, Fig. S6), indicating that task trajectories were substantially more alignable across animals than expected by chance.

We next asked whether the conserved latent geometry across animals was sufficient to support direct transfer of task decoding between animals. To test this, we trained animal-specific CEBRA models independently for each rat and evaluated whether task decoders trained on data for one animal could decode task epochs in another animal after latent space alignment. Importantly, this procedure did not use shared neuron identities across rats; transfer depended only on the shared organization of latent task representations. Following orthogonal Procrustes alignment, decoding accuracy was significantly greater than that of shuffled controls for all train/test comparisons, including A→A, A→B, and B→B transfers (paired two-sided t-tests, all p < 10^-15^; Fig. 4f). Mean decoding accuracy across animals was nearly identical across comparison types (A→A = 0.826, A→B = 0.819, B→B = 0.798) and did not differ significantly from decoding performance when train and test used data for the same rat (two-sided t-tests, all p > 0.25). Of note, these analyses used all recorded neurons from each session rather than only neurons shared across sessions, because comparisons across animals do not require cell registration. Consequently, the number of neurons contributing to the latent representations was substantially larger than in the decoding analyses across sessions for a given animal, shown in Fig. 2, which were restricted to neurons tracked across sessions.

These results demonstrate that task representations generalized across animals to the same extent that they generalized across repeated sessions for an individual animal, indicating that the underlying latent geometry was conserved despite both contextual remapping and inter-animal variability.

Importantly, the models used for decoding across animals were trained independently for each animal and were not optimized to maximize similarity across animals. Successful transfer across animals emerged despite the absence of any explicit optimization objective across animals. Together, these findings demonstrate that hippocampal latent representations share a conserved task related organization across both animals and environments.

## Discussion

Our results show that hippocampal population activity associated with a conditioning task exhibits a conserved organization across spatial contexts and across individuals. Although spatial representations in CA1 remap between environments, the relationships among task related population states were preserved across environments for each animal, and across animals. Notably, conserved geometry was observed not only across animals within the same environment, but also across animals experiencing different environments, indicating that the shared relational structure of task related population activity was maintained despite both contextual remapping and individual variability. These findings suggest that task relevant information is preserved through a stable organization of population level activity, potentially enabling learned behaviors to generalize despite changes in contextual representations.

### Task representations are conserved across environments

A central finding of this study is that the relational structure of task related hippocampal population activity was conserved across distinct spatial environments even as spatial representations reorganized. This dissociation between spatial remapping and preserved task structure suggests that task related information is not reducible to spatial coding but instead emerges from stable population level organization across contexts. Our findings support frameworks proposing that hippocampal representations encode relational task structure in addition to physical space^29^. In this view, spatial and task related information coexist within a shared representational neural space, allowing task structure to remain stable even as contextual features change. Consistent with this idea, hippocampal population activity preserved the relationships among task related neural states across environments despite remapping of spatial representations.

Importantly, the conditioning task used here was explicitly designed to remain identical across contexts, allowing us to isolate generalization of task structure from contextual discrimination. Under these conditions, hippocampal population activity preserved task related organization across environments, suggesting that the hippocampus can maintain task relevant information despite changes in contextual information when such separation is behaviorally appropriate^30–32,55,56^.

### Task representations are conserved across animals

Unexpectedly, our results showed that the relational organization of task related hippocampal population activity was conserved across animals. This conservation occurred despite variability in hippocampal activity across individuals, including differences in spatial tuning, remapping, and the absence of correspondence at the single neuron level. This conservation extended beyond overall similarity in activity patterns and instead involved the preservation of latent task geometry across individual animals.

Notably, the similarity across animals was comparable to the similarity observed across sessions within individual animals. This unexpected result suggests that hippocampal task representations follow shared organizational principles across animals rather than being entirely idiosyncratic to individual experience. Consistent with this interpretation, task decoders trained in one animal successfully generalized to another animal after latent space alignment despite the absence of shared neuron identities. Notably, successful transfer was observed even between animals experiencing different environments, indicating that the conserved relational structure was maintained across both contextual remapping and inter-animal variability.

Conservation across animals is particularly notable given the hippocampus’s established role in episodic memory and its sensitivity to individual experience^33–37^. Prior demonstrations of conserved neural population dynamics across animals have largely involved stereotyped motor or motivational behaviors^38–48^. In contrast, hippocampally dependent tasks such as the conditioning task implemented and studied here involve flexible cognitive processes shaped by learning and context^49–52^. The observation that task related hippocampal population structure was conserved across animals therefore suggests that flexible cognitive functions may also rely on shared organizational principles.

## Conclusion

Together, our findings indicate that the hippocampus supports behavioral generalization through a conserved low-dimensional organization of task related population activity across contexts and individuals. The ability to transfer task decoding across animals further suggests that shared relational structure, rather than specific neural identities, may provide a common representational framework for learned behavior. More broadly, these findings suggest that flexible cognition can emerge from conserved population level organization despite substantial variability in underlying neural representations and individual experience.

## Methods

### LEAD CONTACT AND MATERIALS AVAILABILITY

Questions and requests for information should be directed to and will be fulfilled by the Lead Contact, Hannah Wirtshafter. This study did not generate new unique reagents. The data that support the findings of this study are available from the corresponding author.

### EXPERIMENTAL MODEL AND SUBJECT DETAILS

All procedures were performed in accordance with Northwestern Institutional Animal Care and Use Committee and NIH guidelines. Five male Long-Evans rats (275–325g) were sourced from Charles River Laboratories, injected with AAV9-GCaMP8m, implanted with a 2mm GRIN lens, and trained and tested on eyeblink conditioning in two apparatuses (Fig. 1). Animals were individually housed in an animal facility with a 12/12h light/dark cycle.

### METHOD DETAILS

#### GCaMP8m injection, lens implantation, EMG implantation

GCaMP8m injection and lens implantation were completed as reported in Wirtshafter and Disterhoft, 2022, Wirtshafter and Disterhoft, 2023, and Wirtshafter et al., 2026^1–3^. Briefly, rats were anesthetized with isoflurane (induction 4%, maintenance 1-2%) and a craniotomy was performed at stereotaxic coordinates Bregma AP −4.00mm, ML 3.00mm. 0.6µL of GCaMP8m (obtained from AddGene, packaged AAV9 of pGP-AAV-syn-jGCaMP8m-WPRE, lot v175525, titer 1.3E+13GC/mL) was injected over 12 minutes (approximate coordinates Bregma AP −4.00mm, ML 3mm, DV 2.95mm relative to skull); then the syringe was raised 0.2mm and an additional 0.6µL of GCaMP8m was injected. We repeated this process once more and at slightly different coordinates in the craniotomy hole, resulting in 4 total injections.

We then aspirated tissue from the craniotomy site using a vacuum pump and 25 gauge needle. Tissue was aspirated up to and including the horizontal striations of the corpus callosum. A 2mm GRIN lens (obtained from Go!Foton, CLH lens, 2.00mm diameter, 0.448 pitch, working distance 0.30mm, 550nm wavelength) was then inserted into the craniotomy hole and cemented in place using dental acrylic. Animals were given buprenorphine (0.05mg/kg) and 20mL saline, taken off anesthesia, and allowed to recover in a clean cage placed upon a heat pad.

Six to eight weeks after surgery, animals were again anesthetized with isoflurane and checked for GCaMP expression. If expression was seen, baseplates were attached using UV-curing epoxy and dental acrylic. Electrode implantation to record orbicularis oculi electromyographic (EMG) activity occurred in the same surgery as baseplate attachment, as described previously^4,5^. Briefly, a connector containing 5 wires was cemented on the front of the animal’s head; 4 wires were implanted directly above the eye in the surrounding muscle (2 for recording, 2 for electrical stimulation); an additional wire was attached to a connector attached to a ground screw located above the cerebellum, implanted during lens implantation surgery.

#### Behavioral environment and training

Two behavioral apparatuses were used in these experiments: Environment A was a 78.7cm x 50.8cm unscented rectangular enclosure with wire floor and walls and white lighting. Environment B was a 50.1cm x 34.9cm scented (with two dabs of clove essential oil on opposite walls) ovoidal enclosure with white solid floor and walls, and red lighting. Both environments were located at the same spot in the room relative to external cues (see Figs. 1b and S1).

A tether containing a plug to relay the EMG activity and to deliver a shock to the rat’s eye was attached to the eyeblink connector on the rat’s head. The miniature miniscope (Miniscope) was plugged into the cemented baseplate. The Miniscope and EMG cords were attached to a commutator for ease of animal movement.

The CS was a 250ms, 85dB free-field tone (5ms rise-fall time). The US was a 100ms shock directed to the left eye. Shock intensity varied per session per animal and was calibrated, if needed, at the end of a training session for the next session’s training. Shock level was deemed appropriate when a shock was met with a firm shake of the animal’s head.

The trace interval was 500ms and the intertrial interval (ITI) was randomized between 30s and 60s, with a 45s average. EMG signals were amplified (5000×) and filtered (100Hz to 5kHz), then digitized at 3kHz and stored by computer.

A conditioned response (CR) was identified as an increase in integrated EMG activity that exceeded the baseline mean amplitude by more than four standard deviations, sustained for a minimum duration of 15ms. Baseline mean amplitude was calculated during the 500ms preceding CS onset. Additionally, the response had to commence at least 50ms after the conditioned stimulus (CS) onset and before the unconditioned stimulus (US) onset.

The animal’s first exposure to each environment was a 38min exploration session, in which the animal was able to freely move and explore the environment without any conditioning (Figs. 1a, 1b). Animals were then trained in one environment per session, with no more than one session per day, and were considered to have learned the task after reaching criterion (70% CRs in 50 trials) on three consecutive training sessions (termed ‘criterion sessions’) or when the previous four training sessions averaged over 70% (in this instance, only the final three of those four sessions were considered ‘criterion sessions’). Following the last session in environment A, the animal was given an exploratory session in environment B. In the session after that, the animal was tested on eye blink conditioning in environment B, using the same parameters used in environment A.

#### Calcium imaging

Calcium imaging was completed as reported in Wirtshafter and Disterhoft, 2022, Wirtshafter and Disterhoft, 2023, and Wirtshafter et al., 2026^1–3^. Briefly, calcium imaging was done using UCLA V4 Miniscopes^6,7^, assembled with two 3mm diameter, 6mm FL achromat lenses used in the objective module and one 4mm diameter, 10mm FL achromat lens used in the emission module.

### QUANTIFICATION AND STATISTICAL ANALYSIS

Means are presented as mean ± SEM. All analysis code is available at https://github.com/hsw28/ca_imaging and https://github.com/hsw28/Hannahs-CEBRAs.

#### Position and speed analysis

Position was sampled by an overhead camera at 30Hz. Position tracking was done post-recording using DeepLabCut^8^. Position was then converted from pixels to cm and smoothed using a Gaussian filter with a = 2cm. Speed was calculated by taking the Euclidean distance between two spatial coordinates, one just before and one just after the time of interest.

#### Video pre-processing and cell identification

Video pre-processing and cell identification were performed as reported in Wirtshafter and Disterhoft, 2022, Wirtshafter and Disterhoft, 2023, and Wirtshafter et al., 2026^1–3^. In brief, videos were recorded with Miniscope software at 15frames/second. Video processing was done using CIATAH software^9^. Videos were downsampled in space; the mean value of each frame was subtracted from the frame. Each frame was then normalized using a bandpass FFT filter (70-100cycles/pixel) and motion corrected using TurboReg^10^. Videos were then converted to relative fluorescence (dF/F_0_); F_0_ was estimated from the baseline fluorescence of each extracted calcium signal (calcium trace).

Cells were automatically identified using CIATAH^9^ using CNMF-E^11^. Images were filtered with a Gaussian kernel with a = 2 pixels; neuron diameter was set at a pixel size of 8. The threshold for merging neurons was set at a calcium trace correlation of 0.65; neurons were merged if their distances were smaller than 4 pixels and they had both highly correlated spatial shapes (correlation > 0.8) and small temporal correlations (correlation < 0.4).

In vivo calcium imaging involves detecting changes in intracellular calcium levels, which serve as proxies for neuronal activity. Calcium events refer to transient increases in calcium concentration above a threshold level; these crossings putatively correspond to spikes in neuronal firing. These events typically appear as peaks in the data and indicate an active response from the neuron. Calcium traces are continuous recordings of calcium levels over time. Thus, calcium events highlight specific spike-like neuronal activations, while calcium traces provide a full temporal picture of these activations together with baseline activity.

All cells identified using CNMF-E were then scored as neurons or not neurons by a human scorer. Scoring was also done within CIATAH software in a MATLAB GUI. Scoring was done while visualizing, and based on the calcium activity trace, average waveform, a montage of the candidate cell’s Ca2+ events, and a maximum projection of all cells on which the candidate cell was highlighted. The local maxima of the relative fluorescence (ΔF/F_0_) of each identified cell were identified as calcium events.

#### Cell cross registration across sessions and within session

Validation and registration were completed as documented in Wirtshafter and Disterhoft^2^. Briefly, videos underwent five rounds of registration using Turboreg image rotation^10^ with the CIATAH software^9,12^. Background noise, axons, and dendrites were removed using an image binarization threshold of 40% of the images’ maximum value. Cells were matched across sessions using a distance threshold of a maximum of five pixels, with a minimum 2-D correlation coefficient of 0.5. Sessions were aligned to session A(n), the last session in environment A.

Calcium imaging (CaImg) enabled the longitudinal monitoring of the same hippocampal cells over multiple sessions in both environments. For criterion and testing sessions, we observed an average of 459.85±265.31 cells per session per animal, with no significant difference between the number of cells recorded in environment A and environment B (two-tailed t-test, t(24) = -0.56, p>0.05). On average, 132±95 cells were present in both the last criterion session in A and the first testing session in B. This was not a significantly different number of cells that were present, on average, in both the semi-final session in A, session A(n-1), and the final session in A, session A(n) (155±115 cells).

Cross-animal decoding used all recorded neurons from each session (248–910 neurons/session), whereas cross-session decoding analyses were restricted to neurons successfully registered across sessions (30–263 neurons per comparison).

#### Remapping quantification

The rate center was defined as the occupancy normalized location with the maximum number of calcium events while the animal was moving at 5cm/s or faster. Position was binned into 2.5cm square bins. The rate centers in environments A and B, as well as at environment A across days and at environment B across days, were used to align each environment across days, as well as to align environment A to environment B.

#### Use of CEBRA versus alternative methods

We explored several methods for the dimensionality reduction of Ca events before settling on the use of CEBRA for this study. All methods used the calcium data processed as described above. A short summary of each tested method can be found below:

- **Principal Component Analysis (PCA)**^13^: Principal component analysis (PCA) is a statistical method used to reduce the dimensionality of data while retaining as much variability as possible. This linear technique identifies the axes (principal components) in the dataset that maximize variance. The first principal component explains the most variance, the second explains the second most, and so on. Each principal component is a combination of the original features, in this case the activity of individual neurons; these combinations may not always have clear or intuitive meanings. In agreement with previous hippocampal data^14^, PCA required upwards of 15-25 components to capture 95% of the variance of the data. In addition, across trials within a session, across sessions and environments, and across behavioral representations (spatial and task related activity), the manifolds spanned by the largest PCs remained highly similar, with small principal angles in pairwise comparisons. This similarity in the orientation of the leading subspaces suggested that PCA did not distinguish between spatial and behavioral components of the task (Fig. S7).
- **Independent Component Analysis** (**ICA**)^15^: Independent Component Analysis (ICA) is a computational technique used to separate a multivariate signal into additive independent components. ICA operates under the assumption that observed data are linear mixtures of underlying independent sources. It aims to find a linear transformation that maximizes the statistical independence of the new components, each a linear combination of the original features. We found that ICA embeddings were unstable throughout the length of a recording session. In addition, the embeddings did not clearly map onto behavioral states (Fig. S8).
- **Isomap**^16^: Isomap is a nonlinear manifold learning technique that seeks to capture the intrinsic geometric structure of data. Isomap is useful when linear methods like PCA cannot capture the intrinsic structure of the data, as it preserves the pairwise geodesic (curved) distances between points when embedding them in a Euclidean space of reduced dimensionality. Unlike linear methods such as PCA, Isomap can capture nonlinear relationships in the data. Interestingly, when using Isomap only about 5 neural modes were required to account for a 90% to 95% cumulative variance. However, the embedding shape did not relate to any discernible property of either neural data or behavior (Fig. S9). Dimensionality reduction was achieved, but the resulting representations were not interpretable (Fig. S9).
- **MIND**^14,17^: MIND is a decoding method designed for integrating multiple data modalities to predict various features, particularly sensory and motor functions. MIND uses recurrent neural networks whose hidden variables provide a mechanism for remembering previous inputs; this approach is particularly apt for the analysis of time series data such as neural recordings. While MIND was very robust at distinguishing the different environments, it was not equipped to handle relatively short signals separated in time, such as conditioning trials separated by intertrial intervals. Our analyses using MIND resulted in poor and unstable embeddings that could not be analyzed (Fig. S10).

CEBRA^18^ was chosen for this project for its unique ability to capture nonlinear relationships in the data and to create stable embeddings over short and long time periods. Additionally, the components that were separated in the projected latent representations were well isolated from each other and well correlated with observed behaviors. If the CEBRA model uses d latent dimensions, the data is displayed on the surface of a hypersphere embedded in a Euclidean space of dimension (d+1).

#### Use of CEBRA for position decoding

Optimal parameters for decoding the position of each animal from neural activity were determined using an extensive grid search across the CEBRA parameters: learning rate, temperature, and number of iterations. Models created to compare different sessions, such as a model trained on data from session A(n) used to decode data from session B(1), were trained using only cells that were recorded in both sessions. Models were trained on calcium traces of these cells, labeled with the animal’s (X,Y) position. In all cases, 75% of data was used to train the model while 25% of data was held out for performance testing in the session used for training. All models were run 500 times. Optimal embeddings were determined based on the minimum median distance between predicted and true positions. The optimal parameters for each rat were as follows:

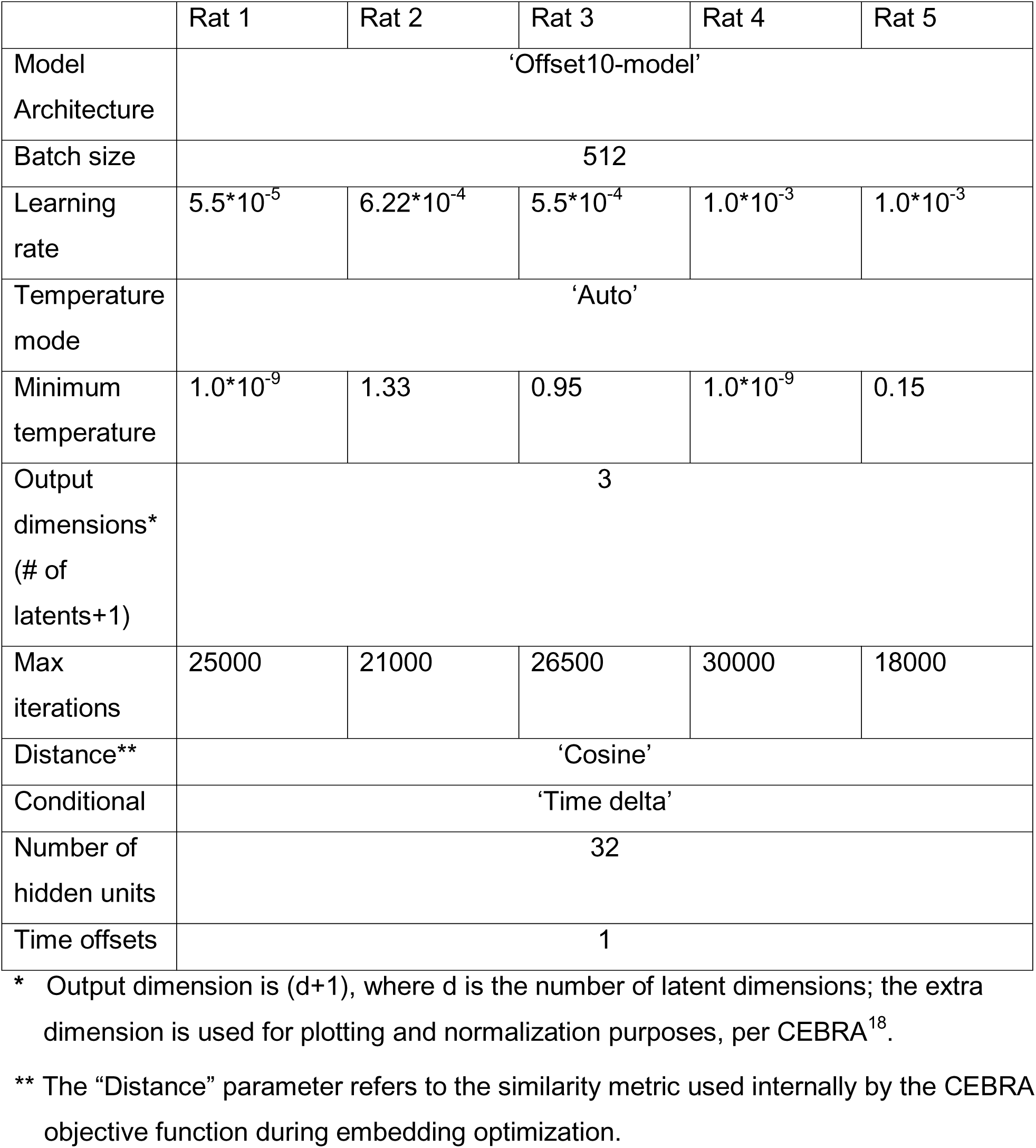

The number of output dimensions was chosen based on the fewest number of dimensions for which all five rat-specific models consistently outperformed shuffled data for both position decoding and temporal decoding during conditioning task execution, as discussed below (Fig. S11-13). To assess chance decoding performance, shuffled controls were generated by randomly permuting the correspondence between latent space data points and position coordinates, thereby preserving the overall distribution of latent neural data while eliminating task related structure. The original CEBRA embeddings were decoded using shuffled labels to generate a control distribution for comparison with decoding of the correctly labeled data.

#### Use of CEBRA for conditioning decoding

As in position decoding, the optimal parameters for temporal decoding during the conditioning task were determined for each animal using an extensive grid search across the CEBRA parameters: learning rate, temperature, and number of iterations.

Models created to compare neural activity across sessions, such as a model trained on data from session A(n) and then used to decode data from session B(1), were trained only on cells that occurred in both sessions. Models were trained on calcium traces of these cells, with labels corresponding to the CSUS bin during which the signal occurred (either 1 out of 2 bins or 1 out of 5 bins). In all cases, 75% of data was used to train the model while 25% of data was held out for performance testing in the session used for training. All models were run 500 times. Optimal embeddings were determined based on the percent of correctly categorized time bins. The optimal parameters for each rat were as follows:

|  | Rat 1 | Rat 2 | Rat 3 | Rat 4 | Rat 5 |
| --- | --- | --- | --- | --- | --- |
| Model | ‘Offset10-model’ |  |  |  |  |
| Architecture |  |  |  |  |  |
| Batch size | 512 |  |  |  |  |
| Learning rate | $5.5 \cdot 10^{-3}$ | $2.5 \cdot 10^{-3}$ | $3.5 \cdot 10^{-3}$ | $7.5 \cdot 10^{-3}$ | $9.5 \cdot 10^{-3}$ |
| Temperature mode | 'Constant' | 'Constant' | 'Auto' | 'Constant' | 'Constant' |
| Minimum temperature | 1.5 | 4 | 1.67 | 1.67 | 2.66 |
| Output dimensions*<br>(# of latents+1) | 3 |  |  |  |  |
| Max iterations | 77000 | 70000 | 20000 | 18000 | 25000 |
| Distance** | 'Euclidean' | 'Euclidean' | 'Cosine' | 'Euclidean' | 'Cosine' |
| Conditional | 'Time delta' |  |  |  |  |
| Number of hidden units | 32 |  |  |  |  |
| Time offsets | 1 |  |  |  |  |

\* Output dimension is (d+1), where d is the number of latent dimensions; the extra dimension is used for plotting and normalization purposes, per CEBRA<sup>18</sup>.
\*\* The "Distance" parameter refers to the similarity metric used internally by the CEBRA objective function during embedding optimization.

The number of output dimensions was chosen based on the fewest number of dimensions for which all five rat-specific models consistently outperformed shuffled data for both position decoding and temporal decoding during conditioning task execution (Fig. S12-14). The parameters listed above were used for decoding into 2 or 5 bins, using 3 output dimensions (2 latent dimensions).

The accuracy of results was computed from the entries in the confusion matrix:

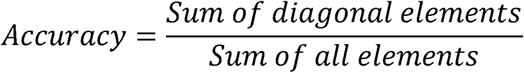

where the diagonal elements correspond to the correct decoding of time bins, while all elements include all decodings, both correct and incorrect.

To assess chance decoding performance, shuffled controls were generated by randomly permuting the correspondence between latent space data points and task labels, thereby preserving the overall distribution of latent neural data while eliminating task related structure. The original CEBRA embeddings were decoded using shuffled labels to generate a control distribution for comparison with decoding of the correctly labeled data.

#### Model consistency

Model consistency was computed using the built-in CEBRA consistency function, which quantifies the similarity between independently trained embeddings. Consistency is assessed by comparing local neighborhood structure across embeddings: for each data point in latent space, the method evaluates whether points that are its nearest neighbors in one embedding remain nearest neighbors in the other embedding. High consistency scores indicate that independently trained models recover similar latent local structure despite differences in initialization.

In the present study, consistency scores were used to compare latent embeddings across environments and across animals. Unlike the geometry-preservation analyses described below, consistency is computed directly from the positions of individual data points within the embeddings and does not require defining task state centroids or computing pairwise distances between centroids. In contrast, geometry-preservation analyses instead quantified whether the relational structure among task state representations is preserved across embeddings.

To quantify consistency between environments and across animals, CEBRA was run 20 times independently for each session; the model achieving the lowest training loss was selected as the representative embedding. Consistency scores were then computed between representative embeddings from different environments and/or different animals using the CEBRA consistency metric. Models for each animal were trained using that animal’s optimal hyperparameters (see above).

To provide a control, chance consistency was evaluated using shuffled data generated by randomly permuting the correspondence between neural data points and task labels. This shuffled data was then used to generate new CEBRA embeddings.

#### Geometry Preservation Analysis

To determine whether task state geometry was preserved when neural activity was remapped across environments, we independently trained CEBRA on session A(n) and session B(1) twenty times for each rat, yielding 20 latent embeddings for A(n) and 20 latent embeddings for B(1). Geometry-preservation scores were then computed for 20 arbitrarily paired A(n)–B(1) embedding comparisons, producing 20 geometry-preservation scores per rat. Because the goal of this analysis was to compare the geometry learned by independently trained latent spaces rather than to evaluate decoder generalization, all available data from each session were used for both CEBRA training and embedding.

Within each embedding, latent coordinates were averaged across all data points corresponding to each of the five CSUS5 time bins, yielding five time bin centroids per embedding. Pairwise Euclidean distances among the five centroids were then computed, producing a 5 × 5 distance matrix containing 10 unique pairwise distances for each embedding. Geometry preservation was quantified as the Spearman correlation between the distance arrays from the paired A(n) and B(1) embeddings. The geometry-preservation value for each rat was defined as the mean of the 20 resulting scores.

To assess significance, we generated a null distribution separately for each rat. For each rat, the observed statistic was a single value: the mean geometry-preservation score across the 20 independently trained CEBRA model runs. To generate the null distribution, we repeated the following procedure 500 times. For each one of the 20 paired CEBRA runs, the labels for the five time bins were randomly permuted for the B(1) session and the resulting geometry-preservation score was computed. The 20 shuffled scores were then averaged, yielding one shuffled mean for that null iteration for that rat. Repeating this procedure 500 times produced a null distribution of 500 shuffled means for that rat. One-sided permutation p-values were calculated as the proportion of rat-specific shuffled means greater than or equal to the observed rat-specific mean; higher correlation values indicate stronger preservation of representational geometry.

The same method was used to compare geometry-preservation scores across rats, except that the distance matrix across rats was computed after averaging all task trajectories for each rat.

#### Embedding Visualization Using Orthogonal Procrustes Alignment

To visualize shared task structure across rats, we generated one representative task trajectory per rat from the CEBRA embeddings of CSUS5 activity. For each rat, embeddings for sessions A(n) and B(1) were first averaged across 20 CEBRA runs to obtain one embedding centroid per time bin for each of these two sessions. Before Procrustes alignment, the resulting centroid coordinates were then z-scored across time bins within each latent dimension, separately for A(n) and B(1). Because the geometry-preservation analysis for individual rats showed that task geometry was conserved between A(n) and B(1), the data representing environment B was orthogonal Procrustes aligned^19,20^ to the data representing environment A for each rat, using corresponding time bins. The aligned B(1) embedding was then averaged with the A(n) embedding to obtain a single representative A/B task trajectory for that rat.

To compare trajectories across rats, each rat’s representative A/B trajectory was orthogonal Procrustes aligned to a common reference rat trajectory. (A single reference rat trajectory was randomly selected only to place all rat trajectories in a common coordinate frame; since Procrustes alignment preserves relative task bin geometry, the choice of reference rat therefore does not define the task structure itself.) Procrustes alignment allowed for rotations, reflections, translations, and uniform scaling while preserving the relative geometry. The aligned trajectories were plotted in a shared coordinate system, with points corresponding to time bins and lines connecting time bins within each trajectory in the appropriate task order.

This visualization was used as a qualitative illustration of the conserved geometry of the task manifold across rats; it complements the quantitative analysis of distance correlations across rats.

#### Decoding Across Animals

We performed task state decoding in CEBRA latent space to test whether the task temporal latent structure learned from one animal could support decoding in another animal. As opposed to matching across sessions for a given animal, matching across animals cannot rely on the matching of individual neurons. To circumvent this limitation, CEBRA was trained independently for each rat using that rat’s neural activity. Transfer between animals was evaluated only after the rat specific latent spaces had been aligned; decoder training and testing did not require shared neuron identities.

To account for variability across CEBRA initializations, we independently trained CEBRA 20 times on session A(n) and 20 times on session B(1), using all available neurons for each CEBRA run. We evaluated three train/test session comparisons: A→A, B→B, and A→B. For each comparison, and for each ordered train-rat/test-rat pair, a decoder was trained on the train rat’s CEBRA latent coordinates and evaluated on the test rat’s latent coordinates after alignment into the train rat’s latent coordinate frame.

The CEBRA models used to generate these latent representations were trained on the full available activity for each session; held out partitioning was applied during the downstream alignment and decoder evaluation procedure. For each ordered train/test rat pair, data points in latent space were divided into two non-overlapping subsets (50% of trials in each subset). One subset was used only to estimate the cross-rat alignment, and the other subset was reserved for decoder training and evaluation.

Before alignment, latent coordinates were z-scored independently within each rat. This normalization removed differences in latent coordinate scale and variance across rats, allowing alignment to focus on the relative organization of time bins rather than the absolute magnitude of the embeddings. Within the alignment subset, centroids were computed as mean latent coordinates for each time bin, separately for each train and test rat. Latent spaces were then aligned using orthogonal Procrustes alignment. The goal of the Procrustes transform was to match these time bin means, with the test rat means treated as the source points and the train rat means treated as the target points. The resulting transform was then applied to the reserved subset of test rat latent coordinates.

Unlike the geometry-preservation analyses, which compare time bin centroids, cross-animal decoding was performed on held out latent data points after Procrustes alignment. Time bin centroids were used solely to estimate the alignment, whereas decoder training and testing used the full subset of reserved latent data points.

For these comparisons across animals, we used CEBRA to obtain low-dimensional latent embeddings. However, CEBRA was not used for decoding as CEBRA cannot decode across different neural populations. Instead, decoding was performed using multinomial logistic regression with balanced class weights. A decoder was trained on the reserved latent data points for the train rat, and evaluated on the aligned, reserved latent data points for the test rat. The entire alignment and decoding procedure was repeated across 20 random data partitions, and the resulting decoding accuracies were averaged.

Shuffle controls were computed by randomly permuting the time bin labels for the train rat prior to training the decoder. This preserved the latent data structure and label distribution for the test rat, while destroying the relationship latent between coordinates and time bin identity for the train rat. Shuffle decoding was implemented independently for each data split.

## Acknowledgements

This work was supported by an NIA T32 (T32-AG020506/AG/NIA), an NIA R37 (R37-AG008796/AG/NIA), an NINDS R01 (R01 NS113804/NS/NINDS), and a K99 award (K99 MH135062).

This research was supported in part through the computational resources and staff assistance provided by the Quest high performance computing facility at Northwestern University, which is jointly supported by the Office of the Provost, the Office for Research, and Northwestern University Information Technology.

We would like to thank all members of the Disterhoft lab, especially Mackenzie Kneisly. We would like to thank the following individuals for their assistance with CEBRA: Mackenzie Mathis and Steffen Schneider. Additional thank yous to David Wirtshafter for his feedback.

## Author contributions

Investigation, H.S.W.; Formal Analysis, H.S.W.; Writing – Original Draft, H.S.W.; Writing – Review & Editing, H.S.W., S.A.S., J.F.D.; Funding, H.S.W, J.F.D., Supervision, S.A.S., J.F.D.

## Supplementary figures

**Figure S1.**
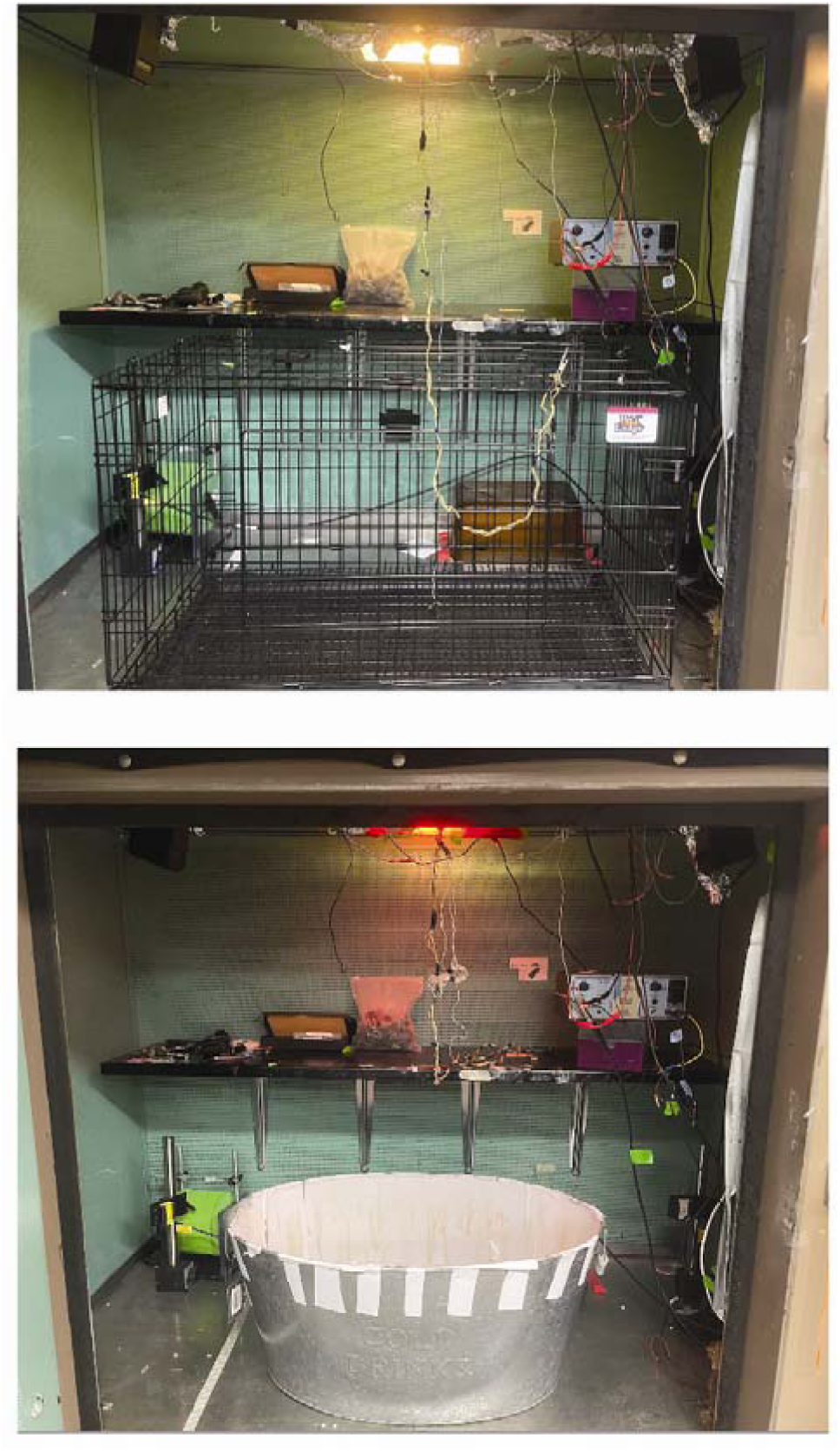
Photos of testing chambers. Photos of testing chambers. Top: Environment A, an unscented rectangular enclosure with wire floor and walls, and white lighting. Bottom: Environment B, a scented ovoidal enclosure with white solid floor and walls, and red lighting. Both environments were located at the same spot in the room relative to external cues. The door to the chamber was closed during testing, to accentuate the distinction between white and red lighting.

**Figure S2.**
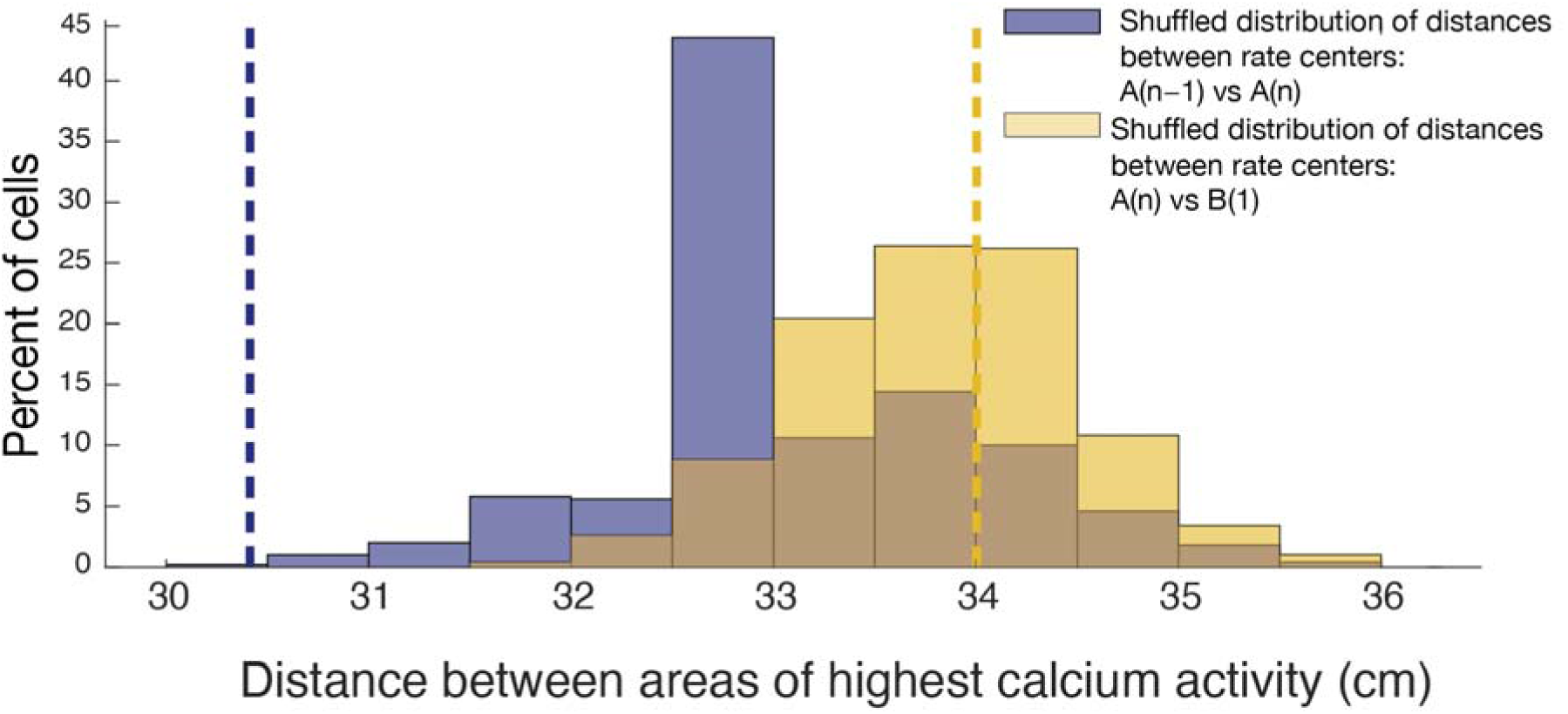
Cells remap across environments, based on comparison to shuffled data. Distribution of median distances between rate centers, the spatial location of peak calcium event rates, after randomly shuffling cell identities 500 times. The observed median distance between consecutive sessions in environment A, dashed blue line, was smaller than nearly all shuffled medians (p = 0.002), indicating preserved spatial correspondence across sessions. In contrast, the observed median distance between environment A(n) and environment B(1), dashed yellow line, fell within the shuffled distribution (p = 0.56), indicating that peak activity locations were no more spatially aligned across environments than expected by chance. The brown shaded area represents the overlap between the two shuffled distributions.

**Figure S3.**
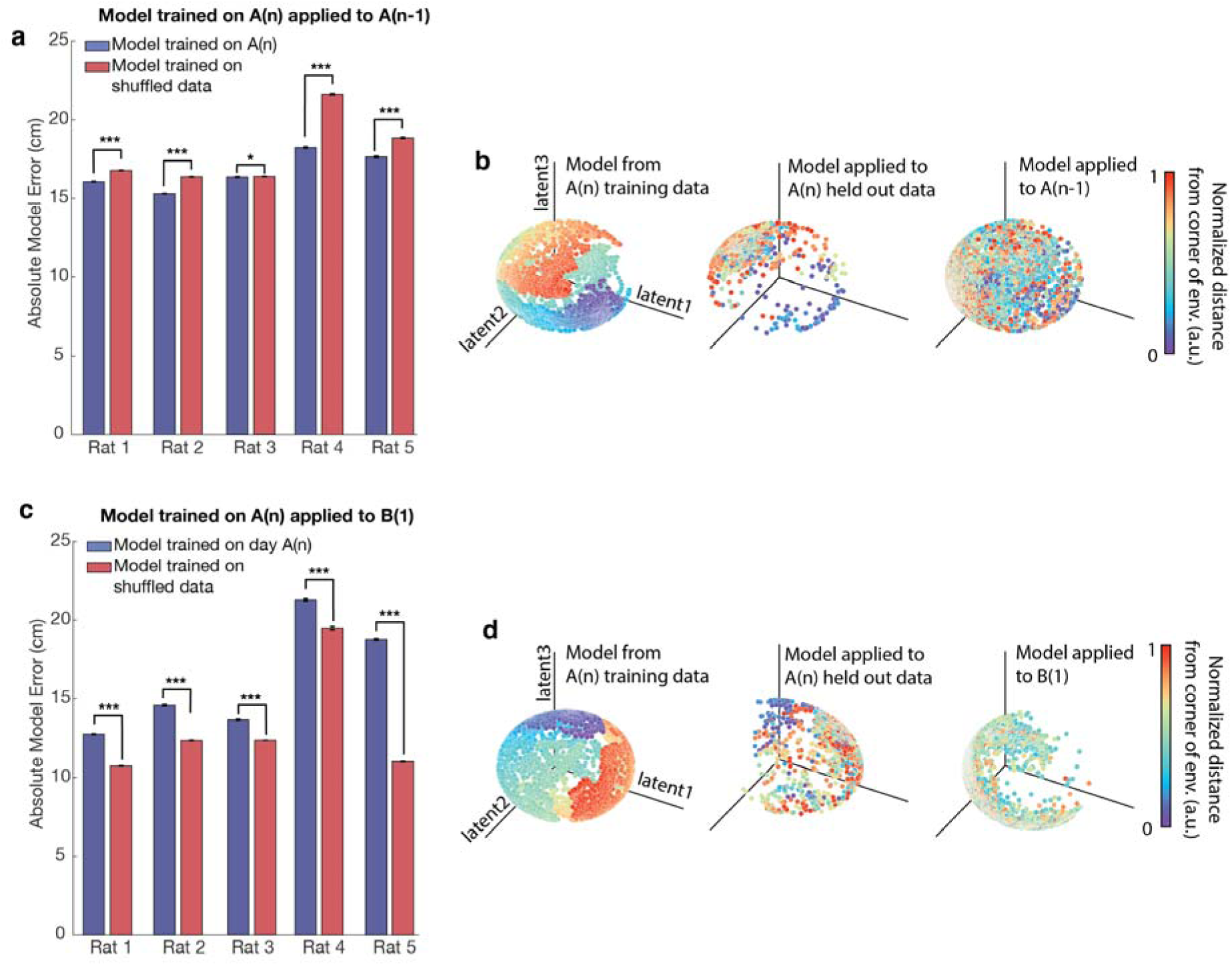
A model trained in environment A can decode positions within environment A but not in environment B. **a.** A model trained on calcium trace and position data from session A(n), using cells present in both A(n) and A(n-1), predicted positions in environment A(n-1) with significantly greater accuracy than that of a model trained on shuffled position data. The model was run 500 times, and the accuracy of predictions was assessed. The model trained on actual data significantly outperformed the shuffled model for all rats (two-sided t-tests: Rat 1: t(998) = -34.3, p = 5.0 × 10^-171^; Rat 2: t(998) = -72.5, p = 0; Rat 3: t(998) = -1.96, p = 0.05; Rat 4: t(998) = - 53.7, p = 2.6 × 10^-299^; Rat 5: t(998) = -20.0, p = 4.1 × 10^-75^). Error bars represent SEM. A single asterisk (*) indicates p ≤ 0.05, and three asterisks (***) indicate p < 10^⁻^□□. **b.** Visualization of model performance for decoding animal position in environment A(n) (Rat 4 example from a single training and embedding CEBRA run). For visualization purposes, distance from the corner of the environment is plotted using normalized values in arbitrary units [a.u.]. Model trained on data from session A(n), using cells present in both A(n) and A(n-1). Left: Model trained on data from 75% of trials in session A(n) (25% of data held out as test data). Middle: Latent representation of held out A(n) data (25%) projected into the latent space learned in one CEBRA run using the complementary 75% of session A(n) task data. Right: Latent representation of data from session A(n−1) projected into the latent space shown on the left, learned in one CEBRA run using 75% of session A(n) task data. **c.** A model trained on calcium trace and position data from session A(n), using cells present in both A(n) and B(1), was applied to environment B(1). The model’s predictions were significantly less accurate than those of a model trained on shuffled position data (two-sided t-tests: Rat 1: t(998) = -84.1, p = 0; Rat 2: t(998) = -18.8, p = 7.4 × 10^-68^; Rat 3: t(998) = -74.7, p = 0; Rat 4: t(998) = -13.1, p = 2.0 × 10^-36^; Rat 5: t(998) = -154.4, p = 0). Error bars represent SEM. Three asterisks (***) indicate p < 10^-35^. **d.** Visualization of model performance for decoding animal position (Rat 4 example from a single training and embedding CEBRA run). For visualization purposes, distance from the corner of the environment is plotted using normalized values in arbitrary units [a.u.]. Model trained on data from session A(n), using cells present in both A(n) and B(1). Left: Model trained on data from 75% of trials in session A(n) (25% of data held out as test data). Middle: Latent representation of held out A(n) data (25%) projected into the latent space learned in one CEBRA run using the complementary 75% of session A(n) task data. Right: Latent representation of data from session B(1) projected into the latent space shown on the left, learned in one CEBRA run using 75% of session A(n) task data.

**Figure S4.**
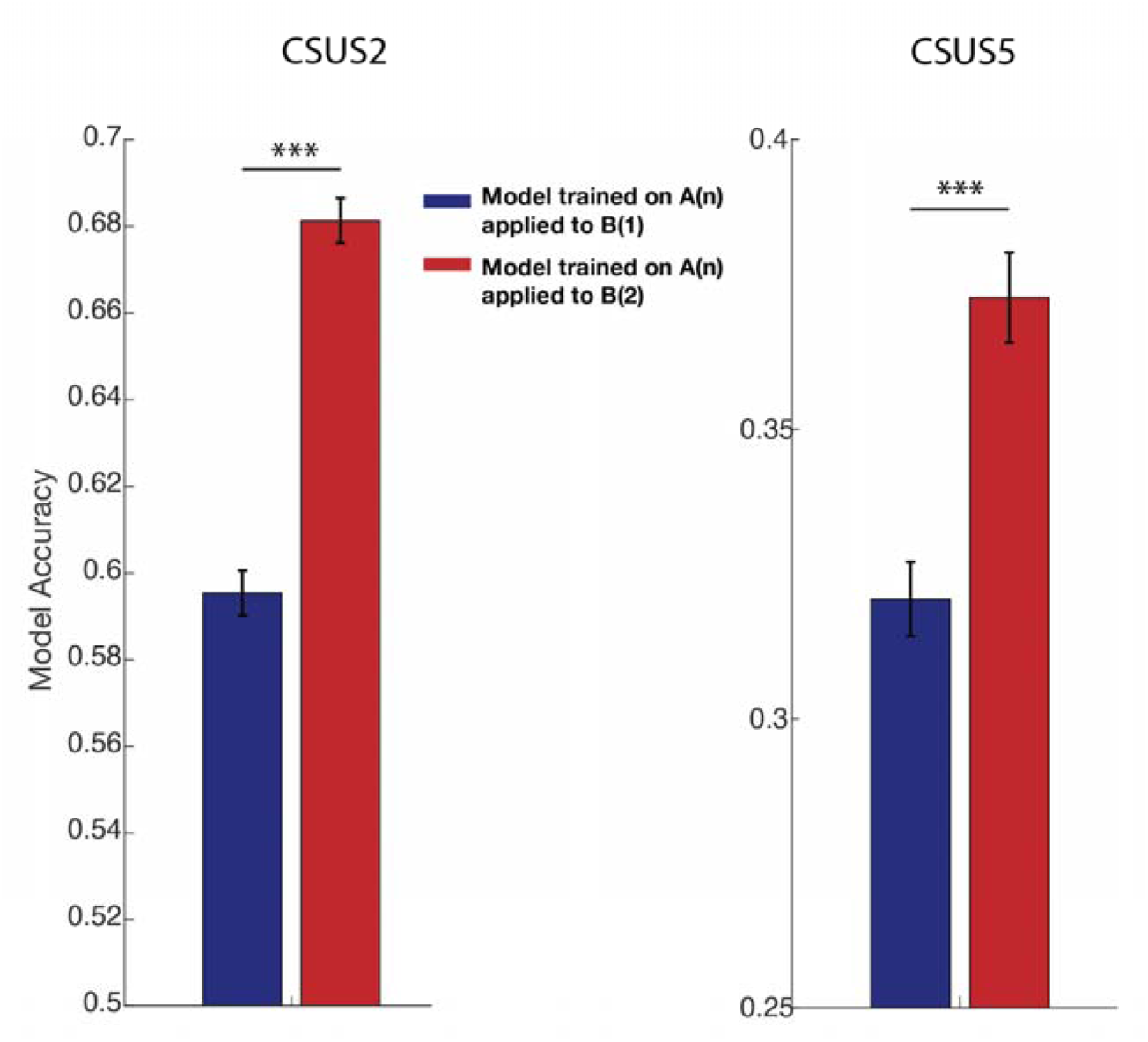
Across environment decoding improves between the first and second exposure to the novel environment for Rat 2. Left: CSUS2 decoding accuracy for models trained on data from session A(n) and tested on data from either session B(1) or session B(2). Right: Same analysis for CSUS5 decoding. In both cases, decoding accuracy improved significantly from B(1) to B(2), indicating stabilization of the transferred latent representation of the tEBC task following initial exposure to the novel environment. Error bars represent SEM. Statistical comparisons were performed using two-sided t-tests (***p < 0.001 for both CSUS2 and CSUS5).

**Figure S5.**
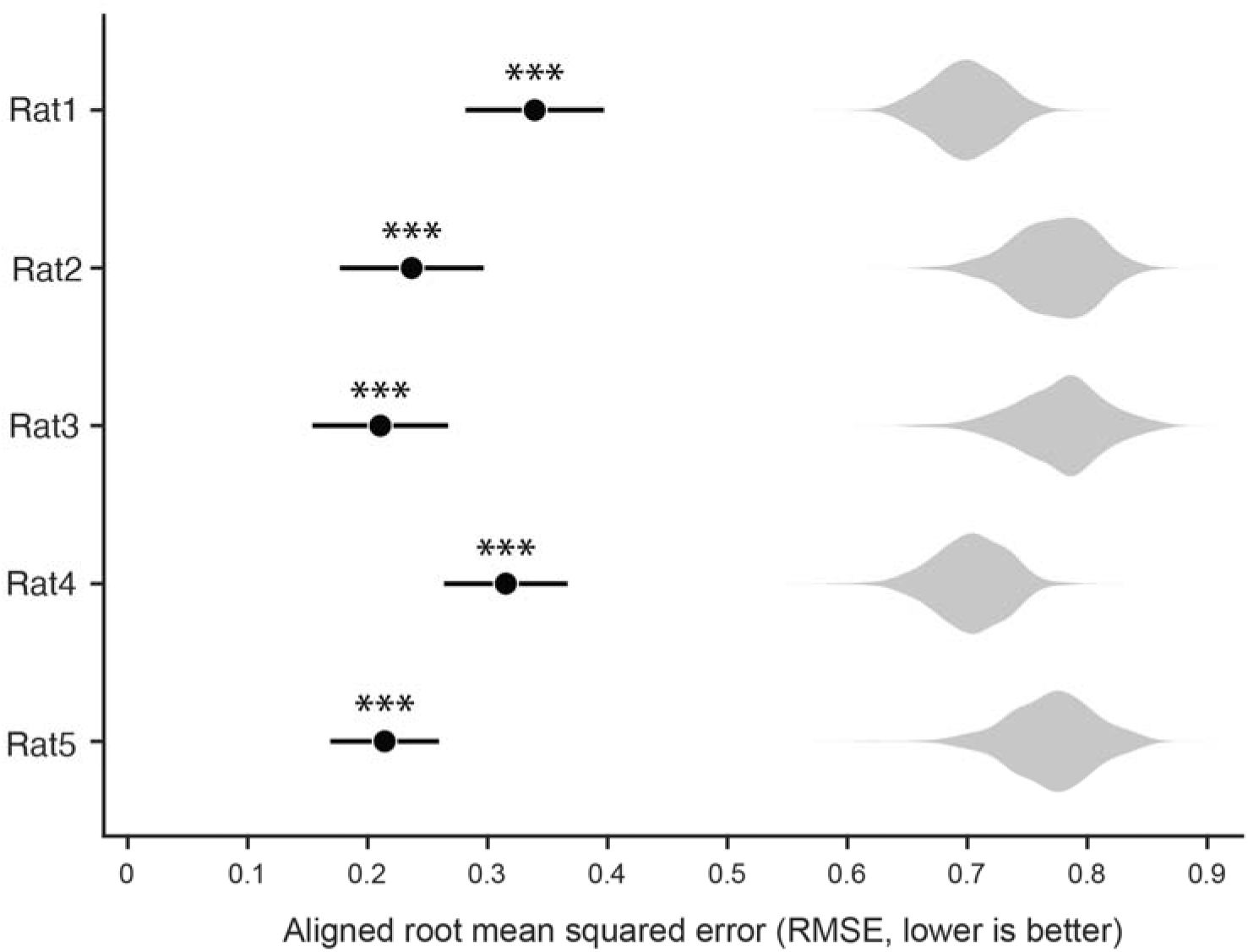
Orthogonal Procrustes alignment error across environments for each rat. To quantify the alignment of task trajectories across environments, latent trajectories from session A(n) and session B(1) were aligned using the orthogonal Procrustes algorithm. Alignment quality was quantified as the aligned root mean squared error (RMSE); lower values indicate better alignment. Black circles indicate the mean aligned RMSE across 20 independent CEBRA runs for each rat; horizontal lines denote ±SEM. Violin plots show control distributions generated by randomly permuting time bin identities in the B(1) trajectories prior to alignment and averaging the resulting RMSE values across runs. Observed alignment errors were significantly lower than expected by chance in all rats (one-sided permutation ***p < 0.002; 500 permutations), indicating that task trajectories were significantly better aligned across environments than expected by chance.

**Figure S6.**
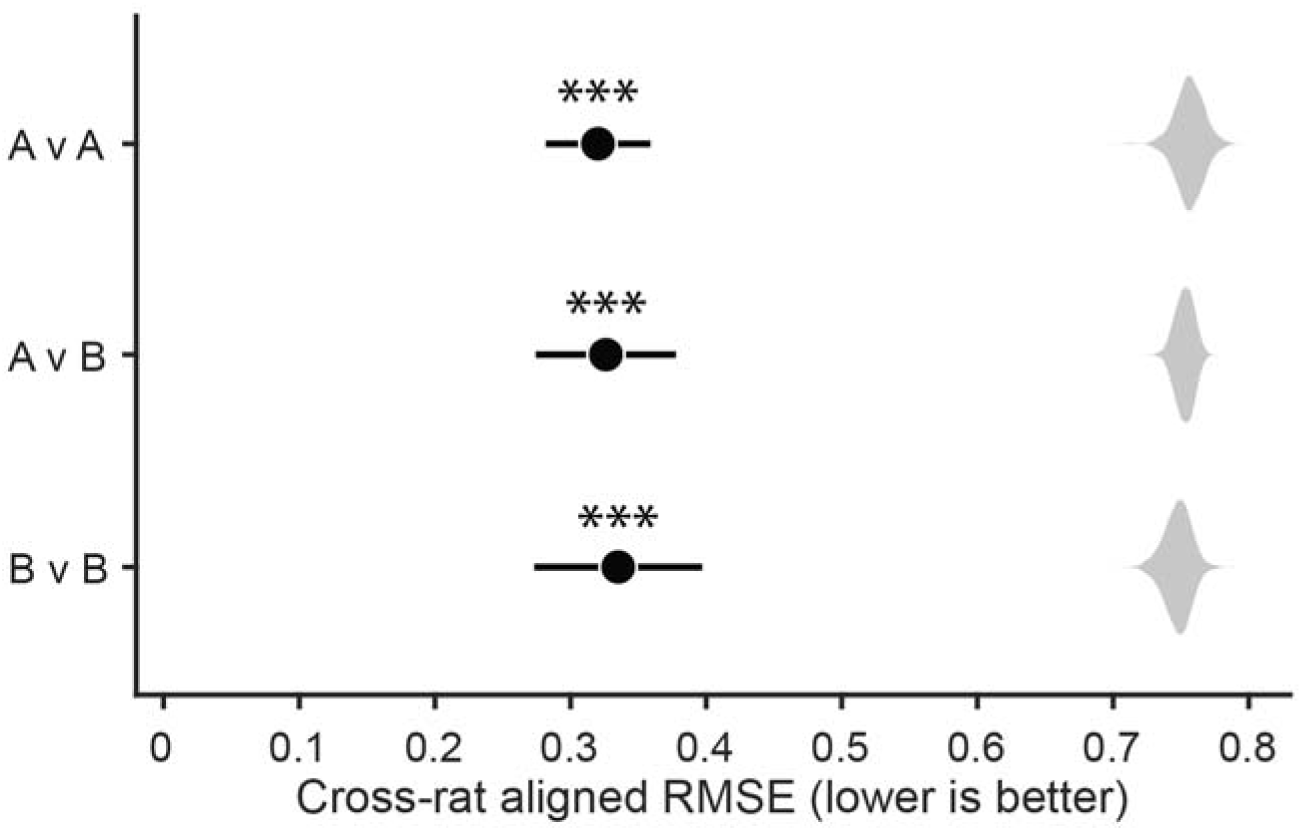
Orthogonal Procrustes alignment error across rats. To quantify alignment of task trajectories across animals, latent trajectories corresponding to the five CSUS5 task states were aligned between pairs of rats using the orthogonal Procrustes algorithm. Alignment quality was quantified as the aligned root mean squared error (RMSE); lower values indicate better alignment. Comparisons were performed using A(n) versus A(n) (A v A), B(1) versus B(1) (B v B), and A(n) versus B(1) (A v B) embeddings across all ten rat pairs. Black circles indicate the mean aligned RMSE across all rat pairs; horizontal lines denote ±SEM. Violin plots show control distributions generated by randomly permuting time bin identities prior to alignment. Observed alignment errors were significantly lower than expected by chance for all comparison types (one-sided permutation ***p < 0.002; 500 permutations), indicating that task trajectories were significantly better aligned across animals than expected by chance, even when the animals were in different environments.

**Figure S7.**
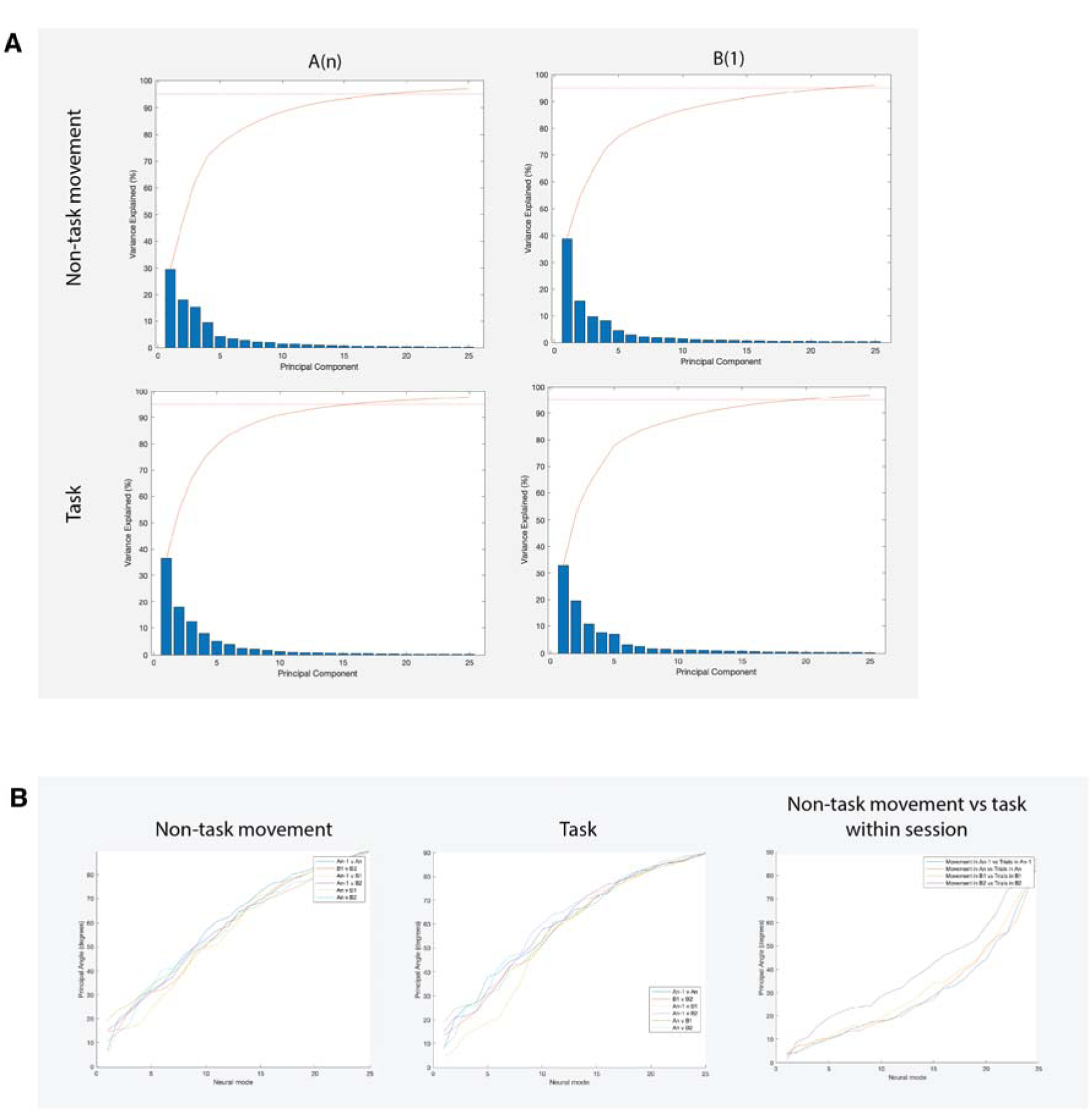
PCA for data from session A(n) and session B(1) for Rat 4. Only cells present in both sessions were used. Principal component analysis (PCA) revealed that approximately 15-25 principal components (PCs) are needed to account for 95% of the variance in the data. When using the complete cell population (not shown), more than 25 PCs are required to account for 95% of the variance. Across and within all sessions and representations (spatial and task), the flat manifolds spanned by the leading PCs are highly similar, as indicated by the small principal angles between them. (A) Eigenvalue distribution (blue bars) and cumulative eigenvalue distribution (variance accounted for, red curve), for both non-task periods (top) and task periods (bottom) in both environments. The black horizontal line shows the 95% threshold. (B) Angles between manifolds for all pairwise comparisons among sessions A(n-1), A(n), B(1), and B(2). Left: comparison of manifolds for activity during non-task periods. Center: comparison of manifolds for activity during task periods. Right: comparison of manifolds for activity during non-task vs task periods within a given session.

**Figure S8.**
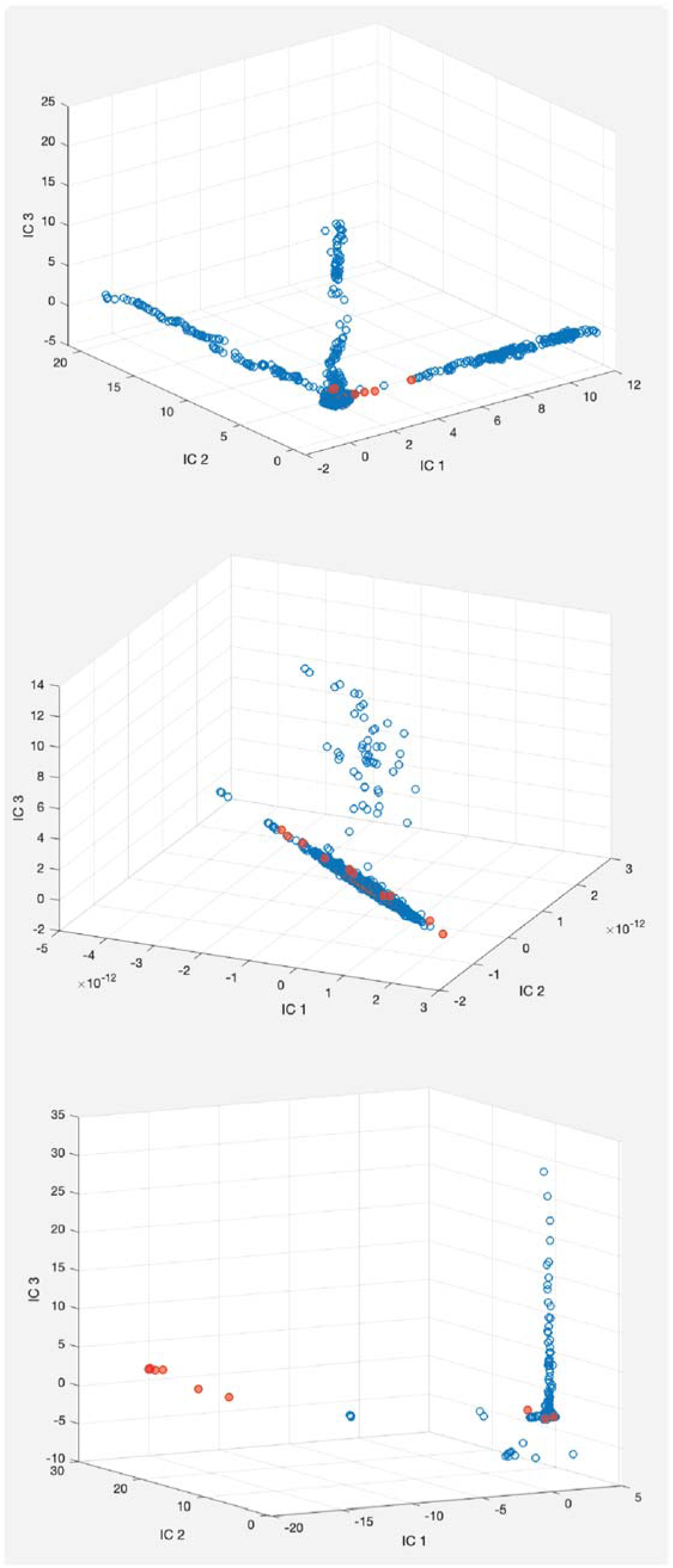
ICA for data from different periods within session A(n) for Rat 4. Top: Independent component analysis (ICA) was performed on data from an entire session, using three independent components (ICs). Blue dots represent non-task times, while red dots represent task times. Middle: ICA computed over the last two-thirds of the same session. Bottom: ICA computed over the second half of the session. Across analyses, the resulting IC structure varied substantially depending on which portion of the session was used. Task-period activity did not form a stable or consistently interpretable organization in the ICA space, indicating that ICA embeddings were sensitive to session partitioning and did not provide a robust representation of task-related activity.

**Figure S9.**
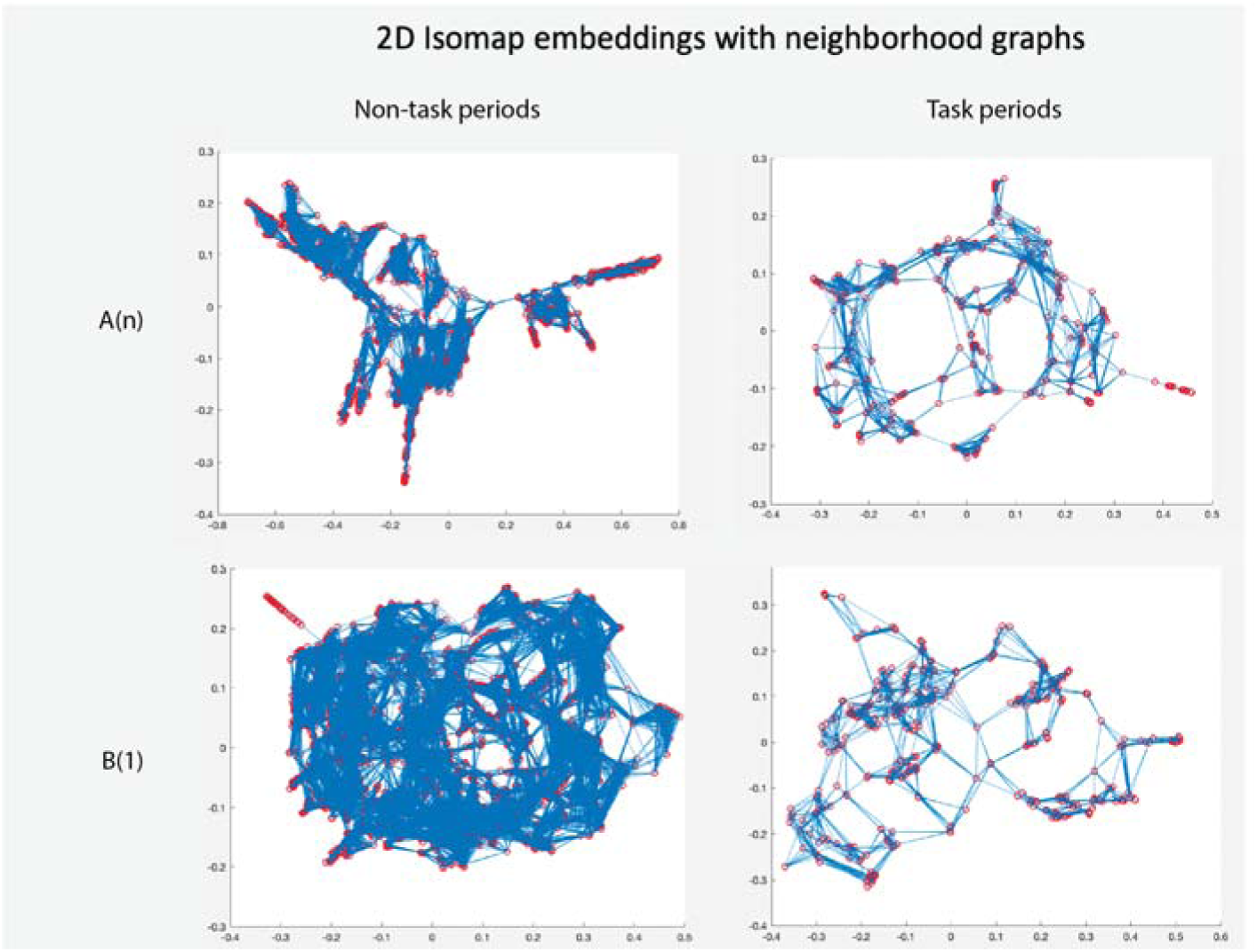
Isomap embeddings for session A(n) and session B(1) for Rat 4. Isomap was computed separately for neural activity recorded during non-task periods (left column) and task periods (right column), using all available data from session A(n) (top row) and session B(1) (bottom row). Red circles indicate individual neural activity samples (nodes), and blue lines indicate the nearest-neighbor connections used by the Isomap algorithm to construct the neighborhood graph. Although Isomap suggested that approximately five neural modes were sufficient to account for 90–95% of the variance, the resulting embeddings did not correspond to any readily interpretable behavioral or neural variables. In particular, embedding structure differed substantially between sessions and did not reveal clear organization related to task state or behavior, limiting the interpretability of the resulting low-dimensional representations.

**Figure S10.**
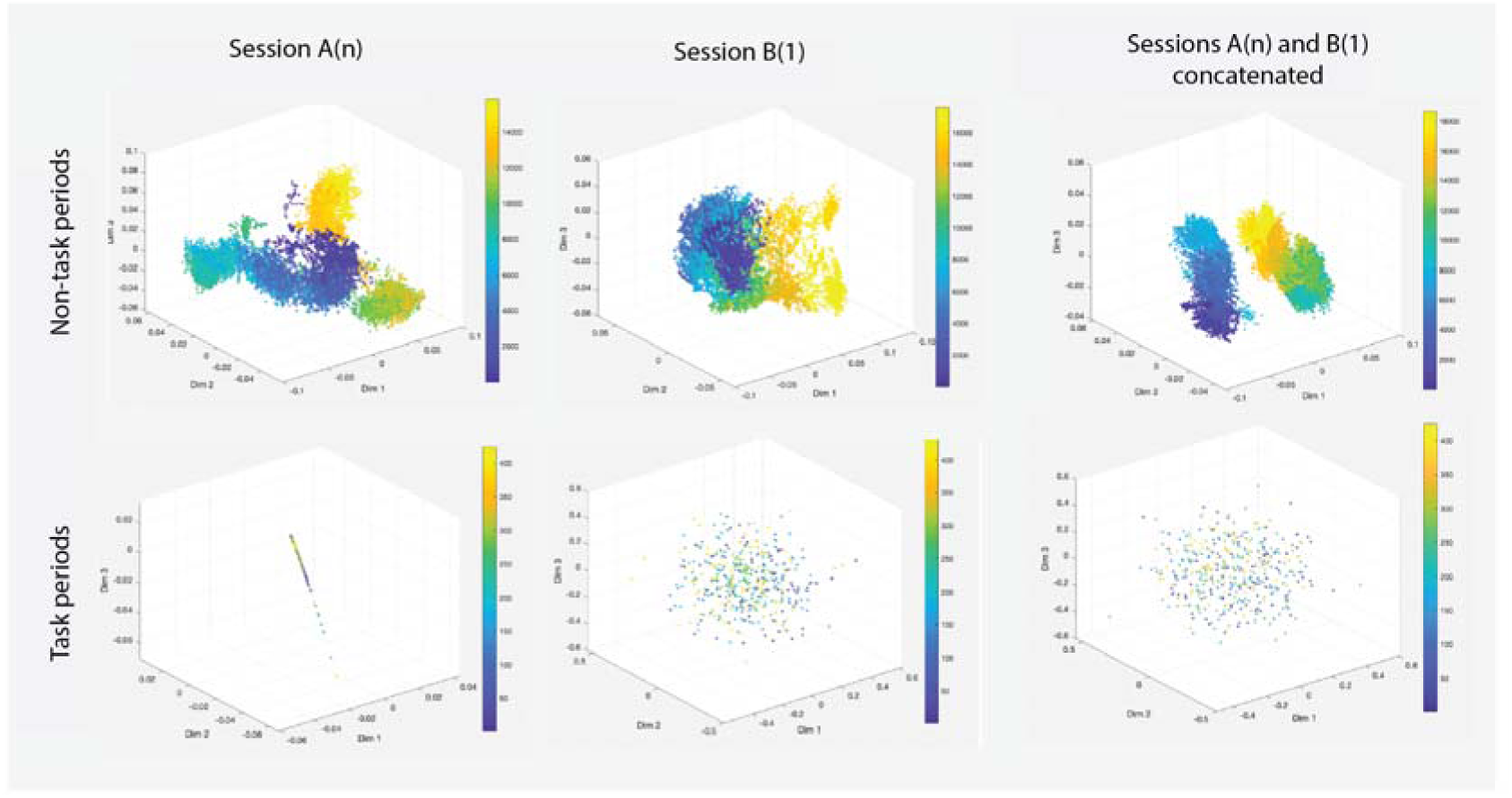
MIND for data from different periods for session A(n) and session B(1) for Rat 4. Top row: MIND embeddings computed from non-task periods in session A(n), session B(1), and the concatenated dataset. Each point represents one sample of neural activity. Colors indicate temporal progression through the recording session (earlier samples shown in dark colors, later samples shown in yellow) and are shown only to aid visualization of the embeddings. The separation between data from sessions A(n) and B(1) in the concatenated embedding (right) reflects remapping between environments. Bottom row: Example MIND embeddings computed from task periods. Task-period embeddings were highly unstable and sensitive to parameter selection. Across repeated analyses, small parameter changes produced qualitatively different embedding structures, ranging from approximately linear arrangements to diffuse clouds.

**Figure S11.**
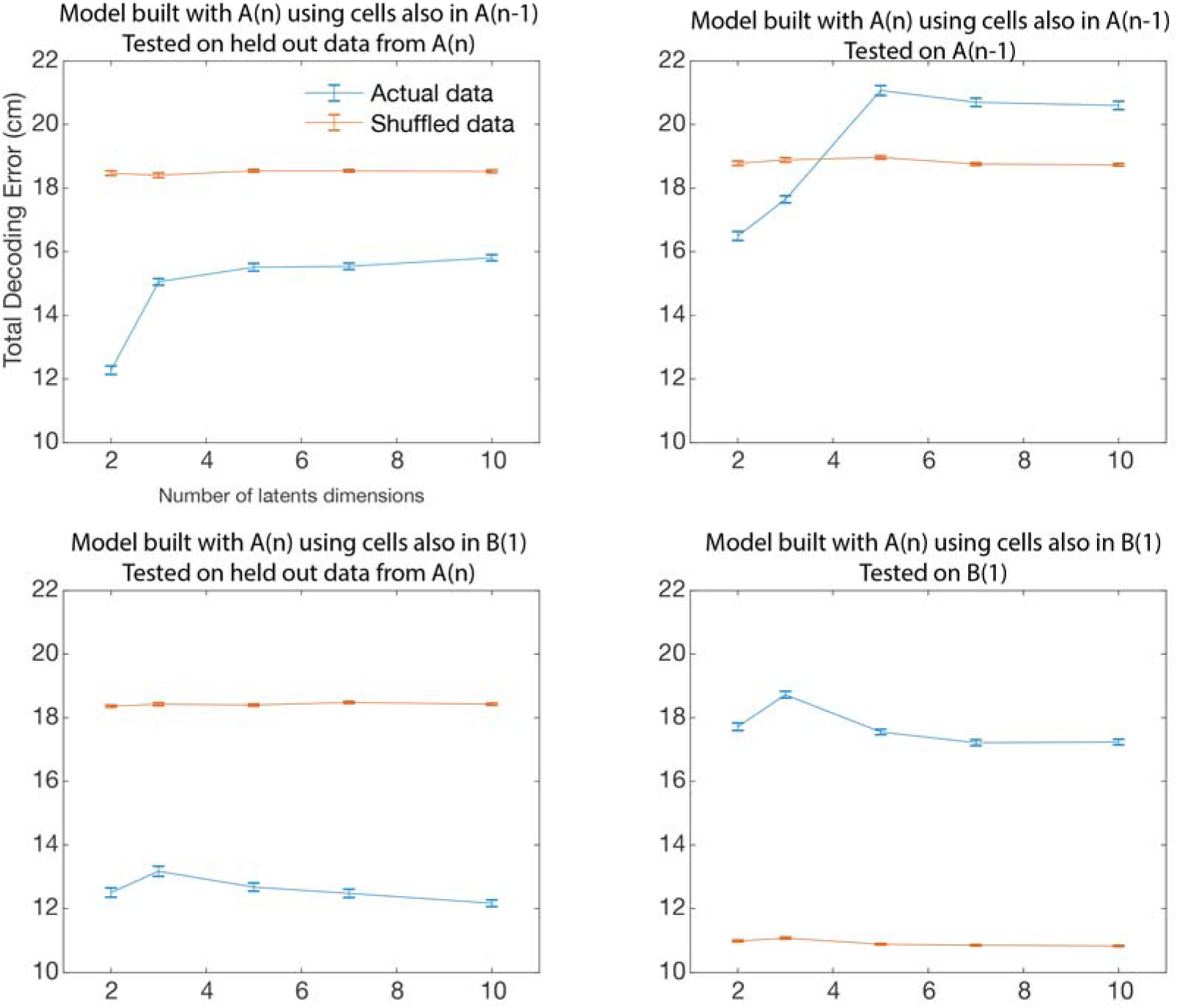
Position decoding error as a function of the number of latent dimensions for Rat 5. Decoding error for position as the number of latent dimensions increases. Left panels: model trained on data from A(n), tested on held out data from A(n). Upper left panel: As the number of latent dimensions increases, the model’s ability to decode held out data from the same session remains stable or decreases. Right panels: model trained on data from A(n), tested on A(n-1) (top) or B(1) (bottom). Performance deteriorates when the model is tested on a different environment (lower right panel). Each model was run 100 times.

**Figure S12.**
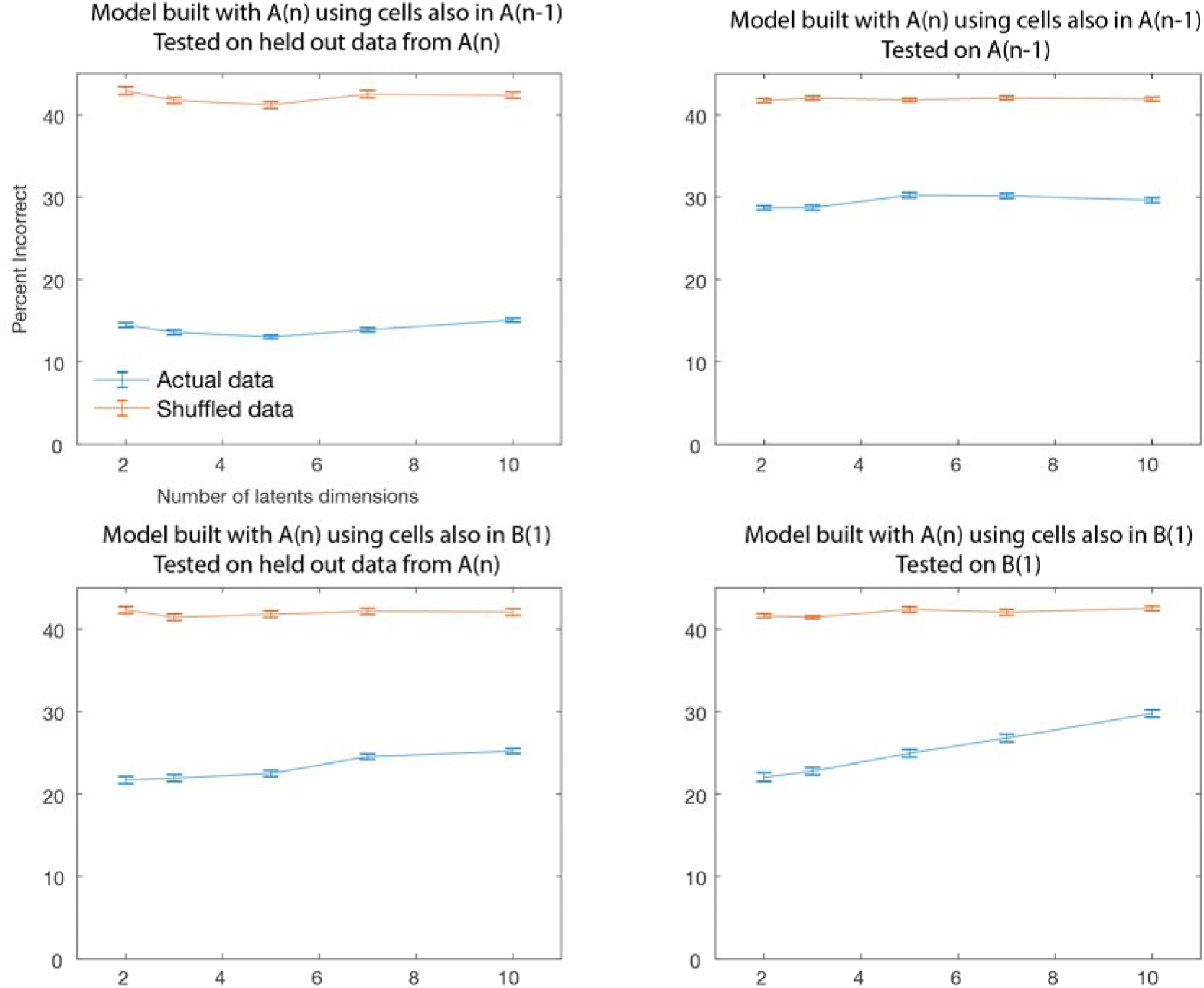
CSUS2 decoding error with increasing number of latent dimensions for Rat 3. Percent of incorrect temporal decoding for the CSUS2 model as the number of latent dimensions increases. Left panels: model trained on data from A(n), tested on held out data from A(n). As the number of latent dimensions increases, the model’s ability to decode held out data from the same environment remains stable or slightly decreases. Right panels: model trained on data from A(n), tested on A(n-1) (top) or B(1) (bottom). Performance remains stable for a different session in the same environment (upper right panel) but deteriorates with increasing number of latent dimensions when the model is tested in a different environment (lower right panel). Each model was run 100 times.

**S13.**
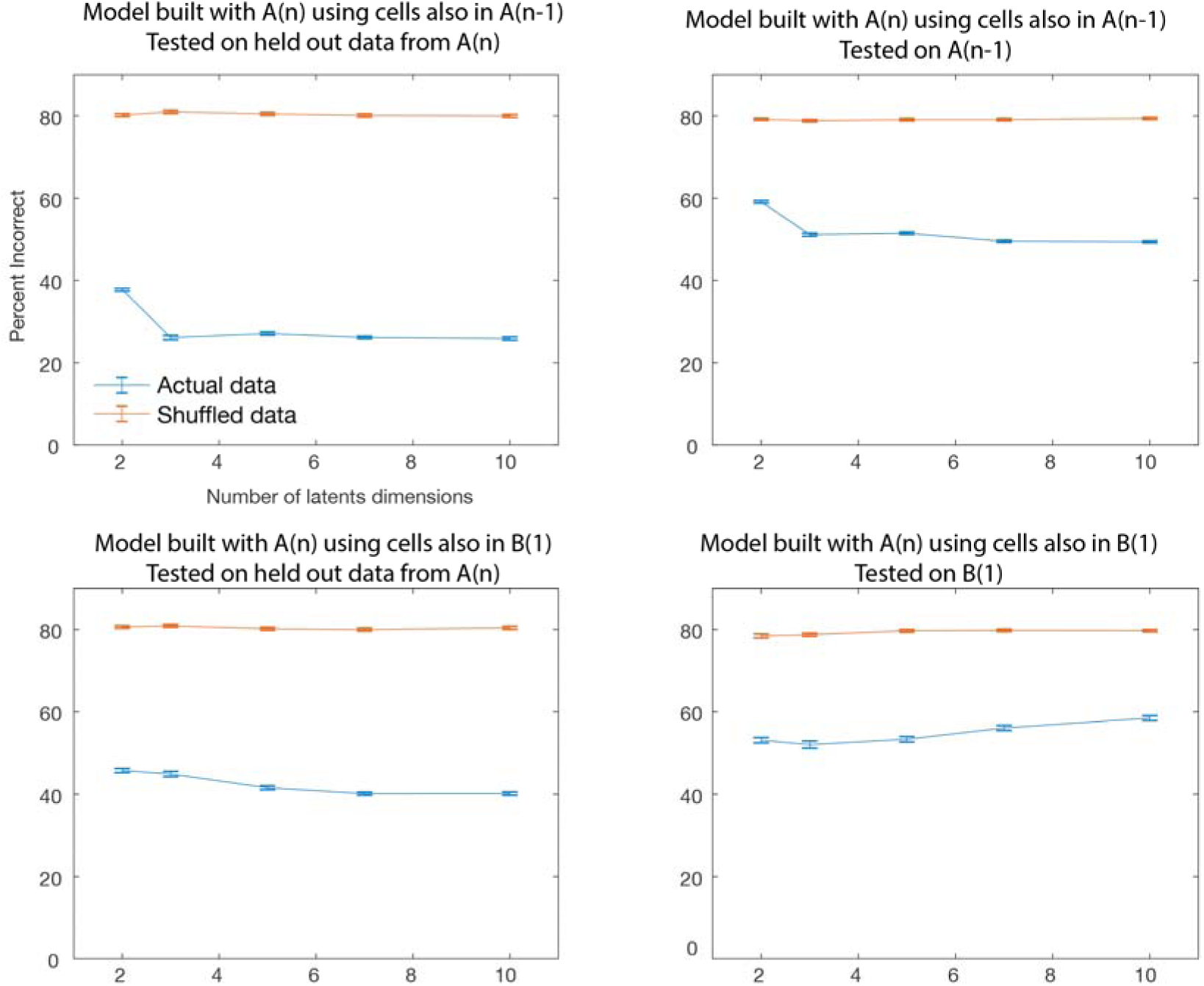
CSUS5 decoding error with increasing number of latent dimensions for Rat 5. Same as Fig. S12, but for CSUS5; the task conditioning period has been divided into five time bins instead of two. The percent of data that is incorrectly decoded is shown as the number of latent dimensions increases. Left panels: model trained on data from A(n), tested on held out data from A(n). As the number of latents increases, the model’s ability to decode held out data from the same environment remains stable or slightly increases. Right panels: model trained on data from A(n), tested on A(n-1) (top) or B(1) (bottom). Performance remains stable for a different session in the same environment (upper right panel) but deteriorates with increasing number of latent dimensions when the model is tested in a different environment (lower right panel). Each model was run 100 times.

